# Characterization of the vertical distribution of plankton and the formation of thin layers in the northern Gulf of Mexico using digital holography

**DOI:** 10.64898/2026.03.29.714992

**Authors:** Gelaysi Moreno, Anvita U. Kerkar, Aditya R. Nayak, Malcolm McFarland, Rubens M. Lopes

## Abstract

The Mississippi River (MR) is the largest source of freshwater and nutrients to the Gulf of Mexico (GoM), strongly influencing stratification, primary production, and plankton organization. The interaction between buoyant plume waters and denser shelf waters in the northern Gulf of Mexico (nGoM) generates sharp density gradients that can promote fine-scale biological aggregation. We investigated how hydrographic structure associated with the MR plume controls the vertical distribution of plankton during May 2017 using an integrated instrumentation suite that included an *in situ* digital holographic imaging system (HOLOCAM) coupled with CTD and optical sensors. Phytoplankton thin layers were repeatedly detected at plume-edge stations within or immediately above a compressed pycnocline formed by bottom-trapped saline wedges. These layers were 1.2–3.5 m thick and exhibited chlorophyll-a concentrations up to threefold higher than background levels. The assemblage was dominated by chain-forming diatoms, particularly *Chaetoceros debilis* and *C. socialis*, whose local abundance maxima coincided with chlorophyll peaks. In contrast, copepods, appendicularians, and other zooplankton were broadly distributed throughout the upper water column and rarely aggregated within the layers. Redundancy analysis indicated that chlorophyll concentration and stratification intensity were primary drivers of community structure across stations. Satellite imagery revealed rapid short-term variability in plume extent, helping explain differences in stratification and thin layer development among sampling days. Our results demonstrate that salt-wedge dynamics at the plume–shelf interface constitute a key physical mechanism governing transient phytoplankton thin layer formation in the nGoM, while zooplankton responses remain weakly coupled at the temporal scales resolved here.

## 1. INTRODUCTION

Planktonic organisms play a crucial role in the biological productivity of the marine environment and, in turn, affect the global carbon cycle and can be indicators of environmental quality and climate variability (Taylor et al. 1986, Sigman et al. 2012, Dyomin et al. 2017). The distribution and variability of phytoplankton biomass and productivity is directly related to the availability of light and submesoscale processes such as fronts, eddies and turbulent mixing that act on the vertical and lateral transport of dissolved nutrients (Hernández-Hernández et al. 2021). Unraveling the interaction of biological and submesoscale physical processes in the formation of phytoplankton patches is crucial to understanding marine ecosystems dynamics (Lévy et al. 2018, Hernández-Hernández et al. 2021).

Vertically compressed and horizontally extensive structures of dense planktonic aggregations within a narrow depth range, known as thin layers, are frequently observed in diverse coastal systems (McManus et al. 2003, Koukaras & Nikolaidis 2004, Penninck et al. 2021) and stratified open ocean waters (Holliday et al. 2003, Sullivan et al. 2010). Thin layers are usually recognized as absolute maxima in vertical fluorescence or beam attenuation profiles (Birch et al. 2008), vertically ranging from a few centimeters to a few meters (Donaghay et al. 1992), while extending to several kilometers in the horizontal direction (Cowles et al. 1998, Churnside & Donaghay 2009), with a temporal duration of hours to several days (Durham & Stocker 2012).

The dynamics of formation and stability of phytoplankton layers are associated with combinations of different physical and biological mechanisms (Frank 1995, Dekshenieks et al. 2001). Potential biological factors include the swimming behavior of phytoplankton (Durham et al. 2009, Sullivan et al. 2010), feeding behavior, population dynamics, reproductive mechanisms, presence of predators and increased local viscosity affecting the displacement of planktonic individuals (Birch et al. 2008, Durham & Stocker 2012).

Several *in situ* optical and acoustic techniques have been developed for the study of small-scale gradients within the water column at high spatial and temporal resolution. The coupling of these sensors to high-resolution imaging systems makes it possible to identify and quantify phyto- and zooplankton abundance in the ocean (Holliday et al. 1998, Benoit-Bird et al. 2010, Sullivan et al. (2005, 2010), Bochdansky et al. 2013, Barua et al. 2024), and investigate the influence of physical, chemical and biological factors on the occurrence, persistence and dissipation of fine-scale aggregations and the vertical distribution of plankton communities in the water column. *In situ* digital holography is one such non-intrusive imaging technique that allows the study of large sample volumes, preserving the positioning of individual particles precisely (Tan & Wang 2013); these key features of the technique have led to its adoption by an increasing number of studies related to particle flow dynamics and biophysical interactions in aquatic ecosystems, such as Nayak et al. (2018a, 2021), Dyomin et al. (2020), and Barua et al. (2023).

The dominant freshwater and terrigenous inputs to the Gulf of Mexico (GoM) can be attributed to the MR and Achtafalaya River (a distributary of the MR) (da Silva and Castelao, 2018; Fournier et al., 2019). These rivers drain in the Texas and Louisiana continental shelf regions of the northern Gulf of Mexico (nGoM), with the MR alone having an annual mean discharge of ∼ 13,000 m^3^/s (Gierach et al., 2013; Fournier et al., 2016), corresponding to >50% of all freshwater discharge into the Gulf. Additionally, freshwater inputs exhibit seasonal trends, peaking in spring (March-June) and being at their lowest in fall (September-November) (Dagg and Breed, 2003; Gierach et al., 2013). Consequently, sea surface salinity (SSS) concentrations also exhibit seasonal variability in the vicinity of the resulting freshwater plume (Fournier et al., 2016). The MR discharges also introduce Colored Dissolved Organic Matter (CDOM) into the nGoM. SSS or CDOM are thus reliable proxies for temporal and spatial tracking or mapping of the MR plume, which can further the characterization of biophysical interactions across scales (Fournier et al., 2016). The complex MR plume dynamics are driven by local topography, wind driven transport and interactions with the Loop Current system in offshore waters (Schiller et al., 2011). The Mississippi River plume is also strongly enriched in nutrients relative to the surrounding continental shelf water, leading to high primary production and influencing plankton distribution and coastal biogeochemistry along the salinity gradient (Lohrenz et al. 1990; Lohrenz et al. 1999; Turner & Rabalais 1994; O’Connor et al., 2016; Liu et al., 2019; Greer et al., 2020).

In this study, observations made on fine-scale plankton community dynamics during a cruise in the nGoM are presented. Specifically, an instrumentation suite equipped with an *in situ* digital holographic imaging system (HOLOCAM), coupled with other optical and acoustic sensors, are used to identify plankton aggregations and characterize their spatial distribution in the nGoM, including specific locations in the vicinity of and within the larger MR plume. The influence of density gradients on plankton vertical distribution and possible thin layer formation mechanisms at different locations of the nGoM was analyzed, to understand the links between plankton dynamics and the physical environment.

## 2. MATERIALS AND METHODS

### 2.1. Sampling and experimental setup

The submersible holographic imaging system (HOLOCAM) employed in this study (Figure 1) uses a 660 nm Nd-YAG laser as the coherent illumination source.

**Fig. 1.**
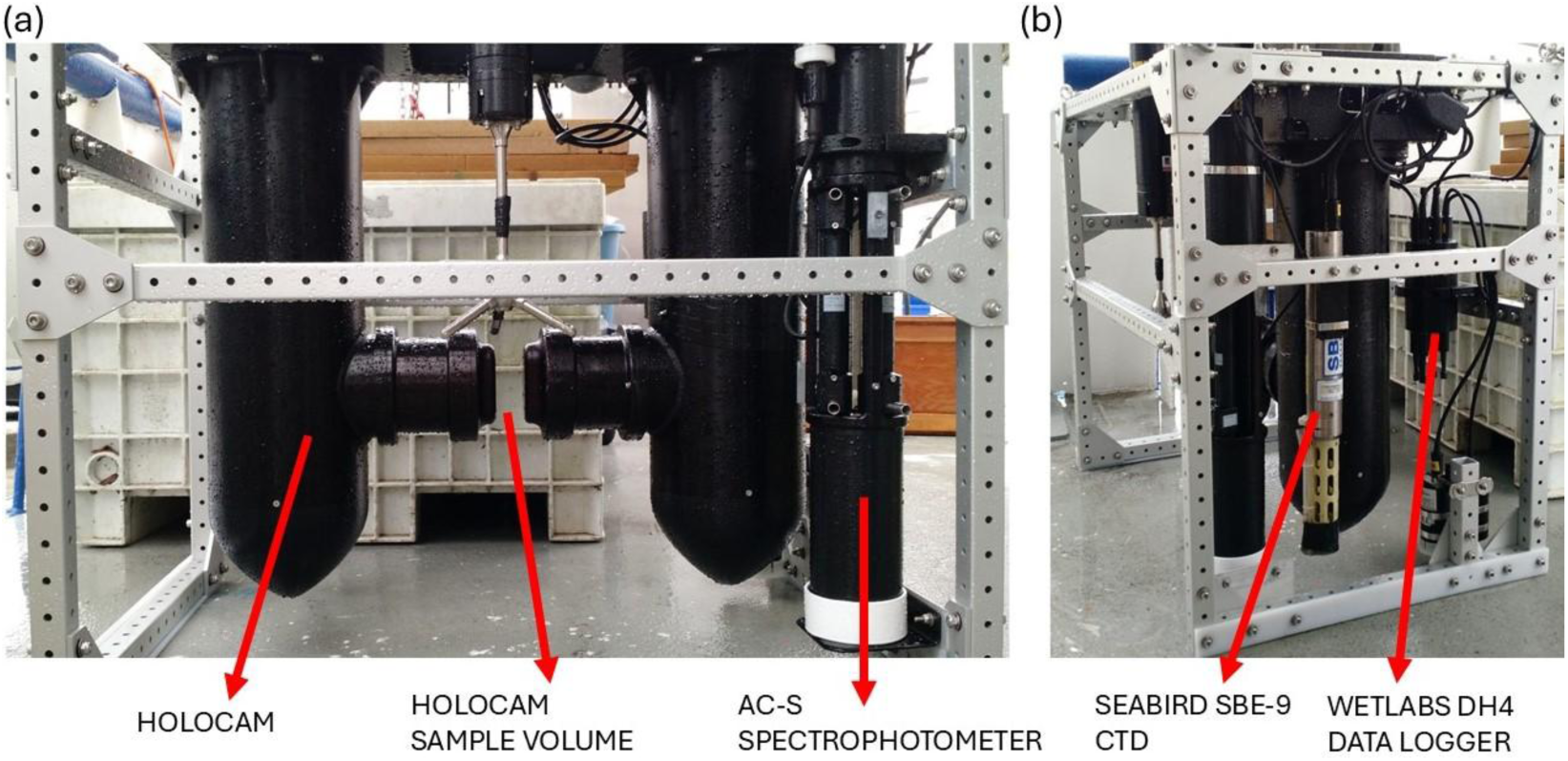
The instrumentation package deployed in the Gulf of Mexico (GoM) with the relevant instruments labelled: (a) Front view of the package showing the HOLOCAM and the ac-s spectrophotometer; and (b) Side view showing the Sea-Bird Electronics 49 FastCAT Conductivity, Temperature and Depth (CTD) sensor and the DH4 data logger.

The resulting holograms are recorded with an Imperx CCD camera (2048 x 2048 pixels) at a resolution of 4.58 μm/pixel, leading to a field of view (FOV) of 9.4 x 9.4 mm and a total sampled volume of 2.65 mL per hologram. Along with the HOLOCAM, the instrumentation suite also included a Sea-Bird Electronics 49 FastCAT Conductivity, Temperature and Depth (CTD) sensor along with a WETLabs ac-s spectrophotometer to obtain simultaneous recordings of salinity, temperature, density (*σ*_*T*_), chlorophyll-a (Chl-*a*) and depth during each vertical profile. For each cast, the instrumentation package was lowered using the ship’s winch, with estimated descent rate of ∼ 0.1 m/s. Only downcasts were considered for analysis of environmental parameters, while both downcasts and upcasts were combined to obtain the vertical distributions of plankton from the holograms. Further details about the instrument package are provided in Nayak et al. (2018a, b).

Sampling was carried out from May 5-12, 2017, between 28.7°N – 29.1°N and - 89.7W – (−90.0°W) south of Louisiana state in the nGoM, including along stations in the vicinity of the MR plume (Figure 2). Some stations were sampled more than once at different times, generating a total of 5 stations and 13 sampling events. Detailed information on sampling time, geographic location, maximum sampled depth, and bottom depth is provided in Table 1.

**Fig. 2.**
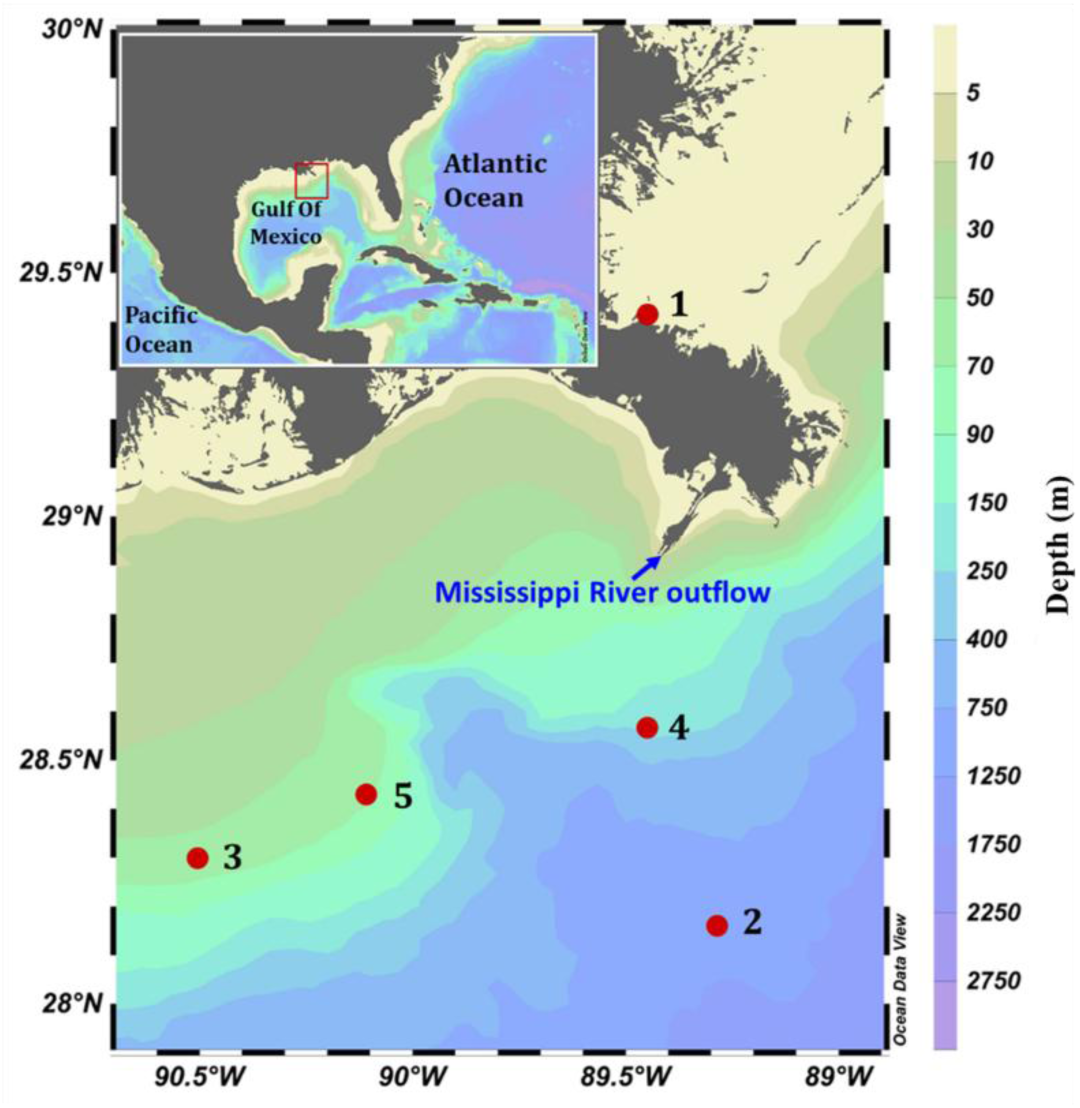
Sampling locations (closed circles) in the northern Gulf of Mexico (nGoM) and an inset view for broader geographic perspective of the study region. The blue arrow marks the Mississippi River (MR) outflow and the color bar indicates bathymetric depth (m).

**Table 1.**
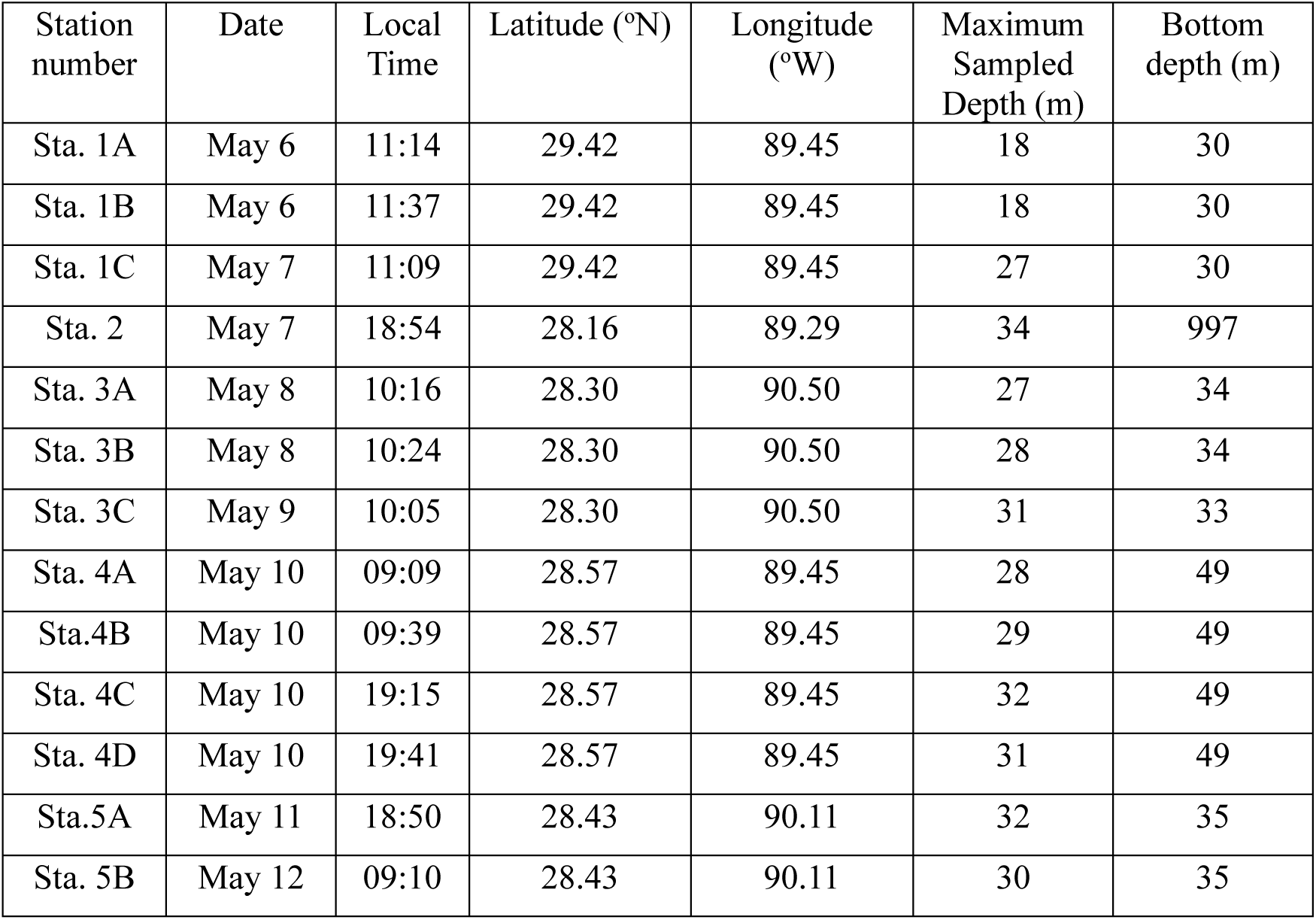
Geographical locations, date, time and depth at each of the sampling stations.

To place the in-situ observations within the regional plume context, satellite-derived products from the Visible Infrared Imaging Radiometer Suite (VIIRS) were analyzed for May 6, 7, and 9, 2017, the only dates during the cruise period with adequate spatial coverage. Level-3 products included chlorophyll-a (Chl-a), enhanced Red-Green-Blue (RGB) composite imagery, and sea surface temperature (SST). Near-surface Chl-a and SST were estimated using SeaDAS software following the algorithms of Hu et al. (2012), O’Reilly et al. (1998), and Brown and Minnett (1999). The enhanced RGB composites were derived from normalized water-leaving radiance at 551, 486, and 443 nm. These products allowed identification of distinct water masses, where brownish tones indicated sediment-laden waters, darker tones represented CDOM-rich waters, and blue regions corresponded to clearer offshore waters with lower particle abundance (Hu et al., 2005).

### 2.2. Data processing

The data processing pipeline to obtain the particle statistics from each hologram can be described in five steps: (1) detection, (2) reconstruction, (3) extraction of relevant features from the images using a convolutional neural network (CNN) (InceptionResNet in our case), (4) dimensionality reduction and (5) data visualization using an interactive labeling tool described by Goulart et al. (2021) (Figure 3).

(1) Given a set of holograms *S* = {*H*_1_, *H*_2_,…, *H*_*n*_}, we estimated a background image (denoted as *I*_*mean*_) from S computing the mean value of each pixel from all holograms. Then we subtracted *I*_*mean*_ from each hologram and an adaptative threshold was applied to the absolute value of the resulting image. To remove undesirable artifacts, we further applied morphological operators such as dilation, erosion and area opening (Dougherty & Lotufo 2003, Lotufo et al. 2023). This process generated a binary image where the remaining connected components were used as a mask to locate the regions of interest (ROIs) in holograms.
(2) Detected ROIs were reconstructed using the angular spectrum algorithm (ASA) (Weng & Zhong 2008) by varying the reconstruction distance in a range of 0-30 mm corresponding to the section of the light path traversing the sample volume relative to the sensor position. Several reconstructions were performed at different planes along the propagation axis in gradual increments of Δ*d* = 0.5 mm. The *Q* focus criterion was applied over the set of reconstructed ROIs (Moreno et al. 2020). The position (Z) matching the maximum value of the *Q* distribution relative to the reconstruction distance was considered representative of the best focus position.
(3-4) The reconstructed images underwent analysis utilizing InceptionResNet for feature extraction. This process yielded a high-dimensional dataset, subsequently reduced to a matrix of similarities among two-dimensional pairs using the t-Distributed Stochastic Neighbor Embedding (t-SNE) technique (Goulart et al. 2021).
(5) The resulting low-dimensional graph visually represents the similarities among the reconstructed images. These images are then plotted, and the resulting points are displayed in an iterative window (Fig. 4) with automatically classified images appropriately labeled, whenever necessary, by an expert in plankton taxonomy.

**Fig. 3.**
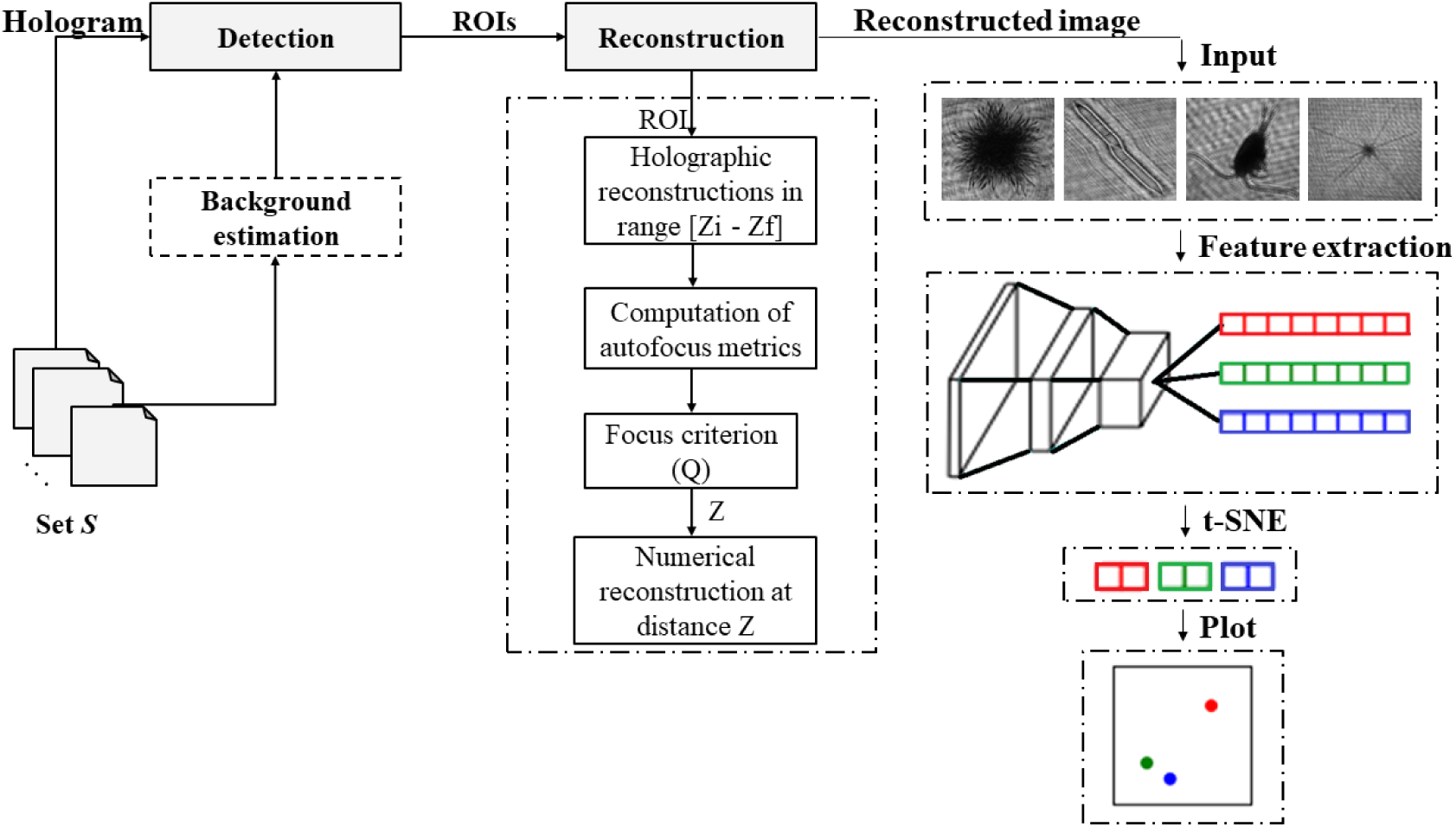
Flow diagram describing hologram processing steps, from particle detection to automatic classification using a convolutional neural network (CNN) and t-Distributed Stochastic Neighbor Embedding (t-SNE).

**Fig. 4.**
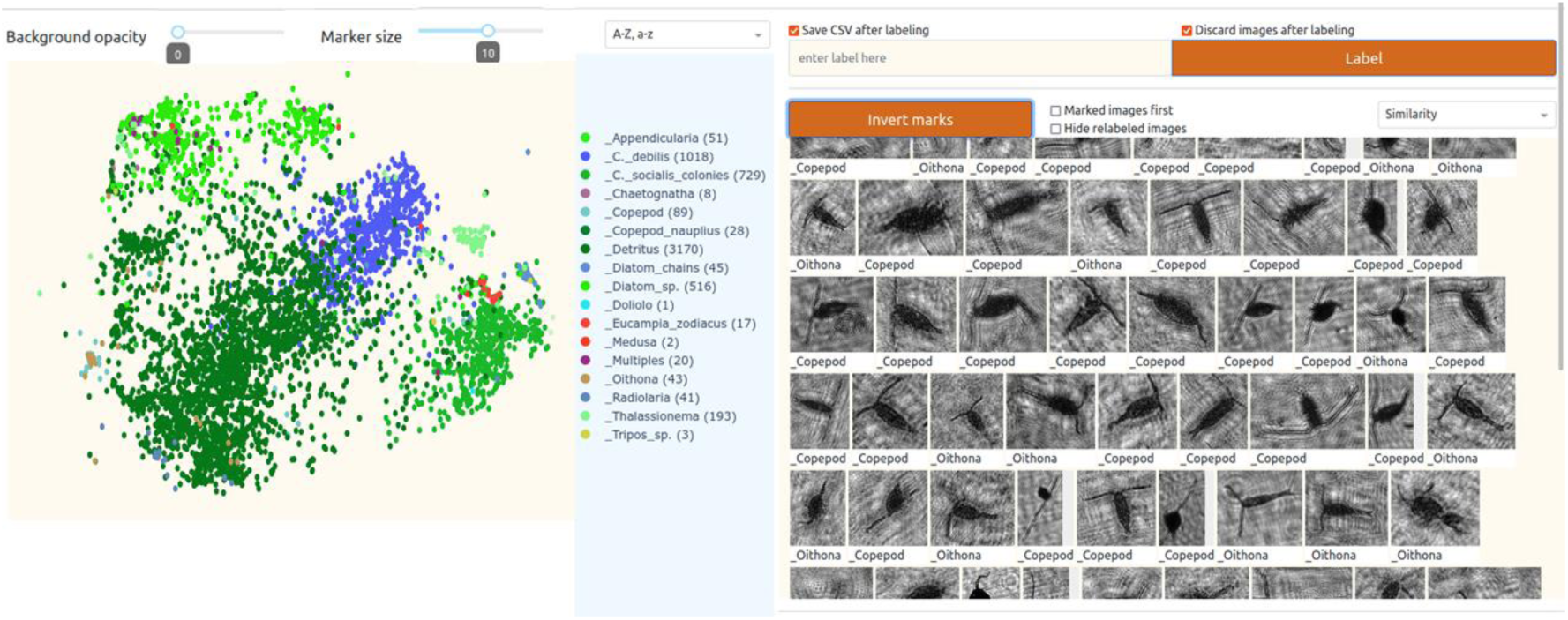
Graphical interface of the Image Labeling Tool (ILT) developed by Goulart et al. (2021), displaying the plotted data points and the automatically sorted images for annotation.

To minimize repeated counts of the same particle in successive frames, we processed every fifth hologram along each cast. Before segmentation, a size threshold of 90 × 90 pixels was applied to exclude smaller objects. Plankton concentrations were calculated in 1 m depth intervals as C_p_ = P/nV, where C_p_ is concentration (particles mL^-1^) in each bin, P the total particles detected in the holograms assigned to the depth bin, n the number of holograms analyzed for that bin, and V the imaged volume per hologram. In all, 15,370 holograms were analyzed – 9,632 from downcasts and 5,738 from upcasts – corresponding to an integrated sampled volume of approximately 40.7 L. Detrital material, including marine snow, fecal pellets and other particles, were not considered in the analysis.

### 2.3. Thin layer identification

Several criteria have been developed to identify which phytoplankton aggregations can be classified as thin layers depending on the detection instrument used, the organisms present in the layer, and the research site (Dekshenieks et al. 2001, Sullivan et al. 2010). The two major criteria to characterize a thin layer are as follows: (1) The layer must have a vertical extent ≤ 5 m thick and (2) the height of the chlorophyll peak must be at least 1.3 times greater than background level, for oligotrophic or mesotrophic waters (Benoit-Bird et al. 2010, McManus et al. 2012) or 3 times higher when considering studies in waters with a higher abundance of planktonic organisms (Durham & Stocker 2012).

Chl-*a* concentration was determined from the absorption values obtained from the ac-s sensor using the specific absorption (a_ph_*; absorption cross-section per mass unit of pigment) line height method at 676 nm, as described in Nardelli and Twardowski (2016). For the calculations, a specific absorption equal to a_ph_*(676) = 0.0108 m^2^/ mg Chl-*a* (Nardelli and Twardowski 2016) was used. Fluctuations in chl-a data were smoothed using a simple 10-point moving average.

The Brunt-Vaisala frequency (*N*), which provides a representation of the local stability of the water column is given by (Pond & Pickard 1983):

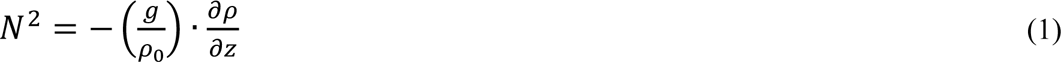

where g is the acceleration due to gravity, *ρ*_0_ is the average density of seawater, *ρ*(*x*, *y*, *z*, *t*) is the local density, and z is the depth. Positive, zero and negative *N*^2^ values correspond to a stable, neutral and unstable water column, respectively. In our case, N varied between 0 – 5 s^-1^ between the different stations, thus showing that the water column was stably stratified.

### 2.4. Statistical analysis

Differences in the abundance of the different phytoplankton groups across the surface, intermediate, and bottom layers were assessed using the non-parametric Kruskal-Wallis test, as the data did not meet the assumptions of normality and homoscedasticity, implemented with the stats package in R (function kruskal.test). Each test included a total of 39 observations (n = 39), equally distributed among the three depth layers (13 samples per layer), resulting in two degrees of freedom (df = 2). When the Kruskal-Wallis test indicated significant differences (p < 0.05), Dunn’s post-hoc multiple comparison tests were conducted using the FSA package (function dunnTest), with p-values adjusted using the Bonferroni correction, adopting the same significance level (α = 0.05).

All satellite image processing and figure production were performed in R v4.3.2 (R Core Team, 2024) using RStudio Posit Team, 2024). Satellite imagery products (chlorophyll-a, enhanced RGB, and sea surface temperature) were imported as raster images and displayed using ggplot2 (Wickham, 2011; 2019) with png, grid, and gridExtra. Multi-panel layouts were created using patchwork (Pedersen, 2020). Station locations were overlaid and labeled to allow spatial comparison across dates and products.

To describe how the most abundant phytoplankton and zooplankton taxa relate to the environmental variables investigated, a Redundancy Analysis (RDA) was performed, following the method of Legendre & Anderson (1999). RDA is a canonical form of Principal Component Analysis (PCA) that selects the linear combination of environmental variables that yields the smallest sum of the total residual variance in the biological data fit. This analysis was conducted considering data from 13 sampling sites, four environmental variables (temperature, salinity, density, and chlorophyll) and 15 dominant phytoplankton and zooplankton taxa. Because taxon abundance exhibited a high deviation from a normal distribution, a Hellinger transformation was applied to stabilize the variance and give less weight to highly abundant species, using the decostand function within the same package. The environmental variables were standardized to have a mean of zero and a standard deviation of one. The RDA was performed using R v4.3.2 (R Core Team, 2024), using the vegan package (Oksanen et al. 2019).

## 3. RESULTS

### 3.1. Environmental conditions

Satellite-derived Chl-*a*, enhanced RGB, and SST maps (Fig. 5) provided a synoptic view of the MR plume during the sampling period and place the hydrographic profiles in a regional context.

**Fig. 5.**
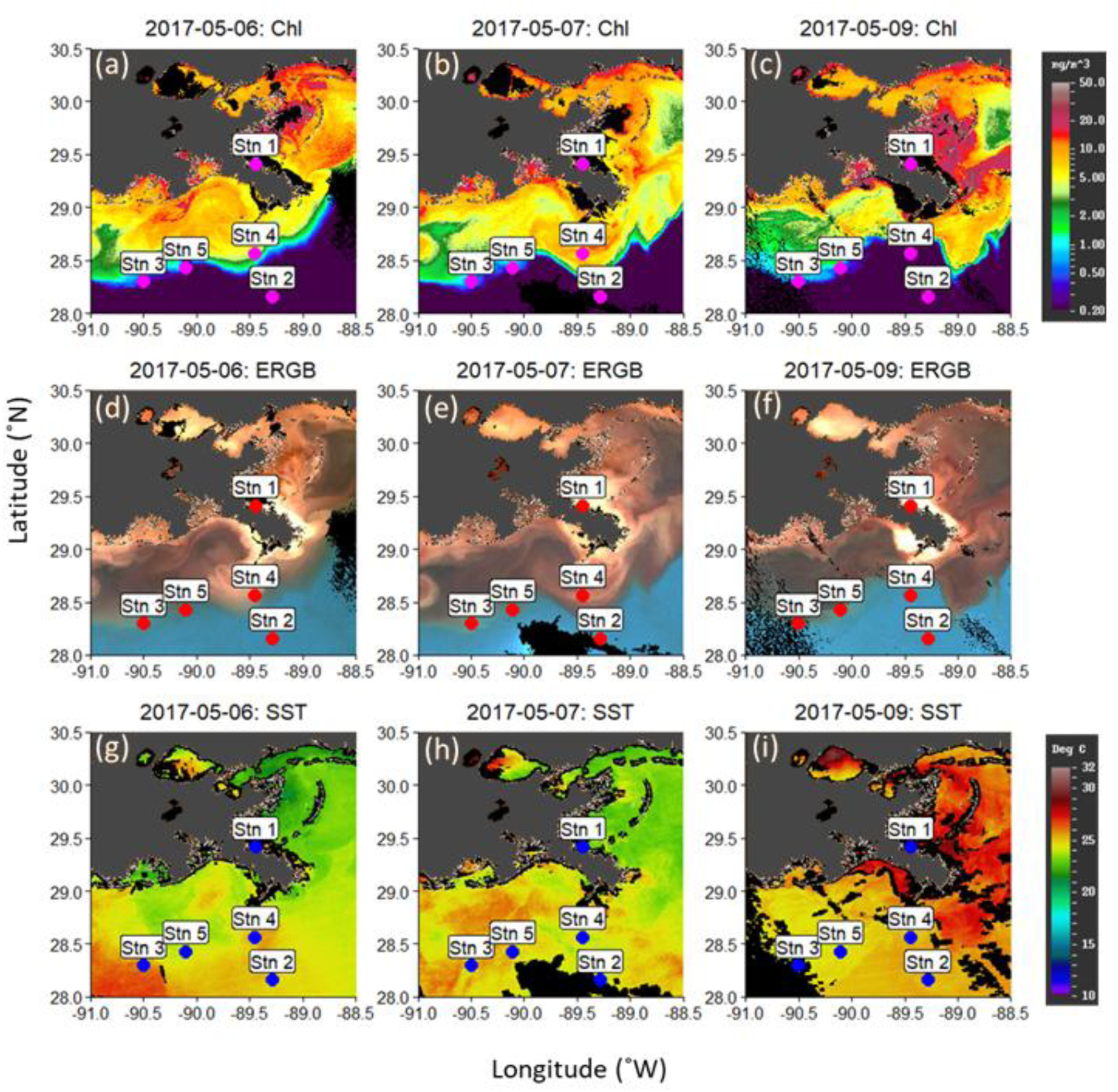
Satellite derived data products for the study region for May 6,7 and 9, 2017, showing (a-c) Chl-*a*; (d-f) composite enhanced RGB image; and (g-i) SST respectively. Sampling locations are denoted by bold circles. These images are adapted from the VIIRS/SNPP Level 3 processed data, publicly available, courtesy of the Optical Oceanography Laboratory at the University of South Florida.

The variations in Chl-*a* and the enhanced RGB maps across the three different days indicated the dynamic nature of the freshwater plume and its extent. Satellite-derived Chl-*a* and RGB composites showed elevated surface pigment concentrations in the region surrounding Station 1, indicating that this site was located within a high Chl-*a* water mass. Stations 3, 4 and 5 were either within or at the edge of high Chl-*a* regions (Fig. 5a-c) depending on the day, which is consistent with the higher phytoplankton abundance in these datasets (see later sections). This spatial gradient observed in Fig. 5 was consistent with the hydrographic structure revealed by the T-S diagrams (Fig. 6) and the vertical salinity profiles (Fig. 7), supporting the classification of stations according to plume influence.

**Fig. 6.**
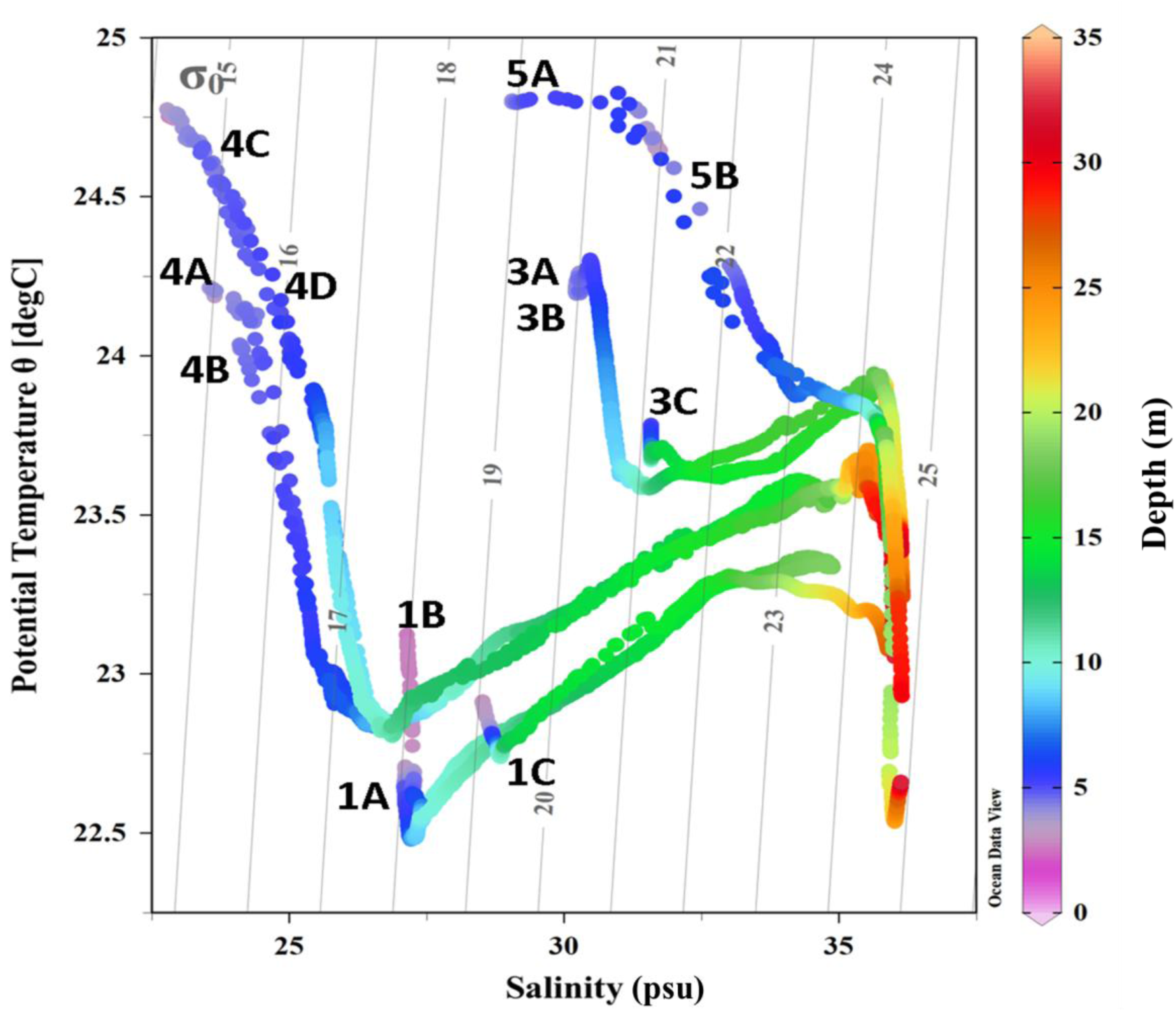
Temperature-salinity (T-S) plots from the CTD data for all the sampled stations, with isopycnals overlaid. Color indicates depth and the respective station numbers are indicated next to each profile.

**Fig. 7.**
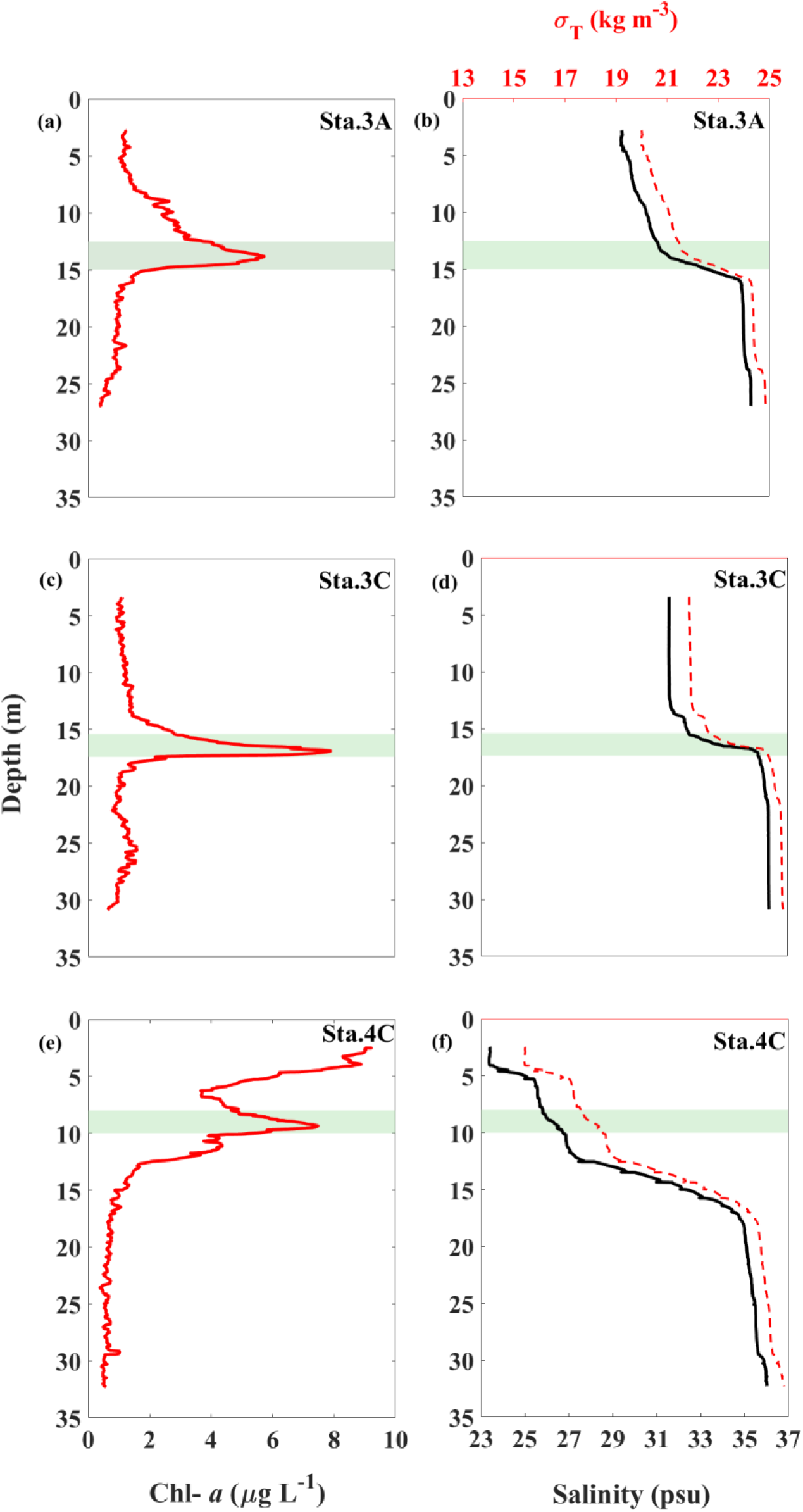
Hydrographic structure of the water column associated with the formation of a salt wedge and phytoplankton thin layers across Stations 3A, 3B and 3C.Vertical profiles of salinity (black), density (*σ*_*T*_; red dashed curves), and chlorophyll-a concentration (Chl-*a*; red) for (a-c) Station 3A, (d-f) Station 3C and (g-i) Station 4C. Shaded region in each profile indicates the depth range of detected thin layers.

Station 2, with a depth of 997 m, was more representative of open ocean conditions with blue waters (low Chl-*a* and particle abundance, and clearer waters). Additionally, Figures 5d-f outlined the sediment plume associated with the MR outflow, indicating Stations 1 and 4 fall within this region and Stations 3 and 5 are in the vicinity, but possibly not within, the main plume. It should be noted that at Stations 4 and 5, our data was collected on May 10, 11 and 12, for which good satellite coverage is not available. Additionally, the SST data revealed high short-term variability, including a ∼ 3-4 °C local increase in some regions between May 7 and 9 (Figure 5g-i).

Fig. 5 revealed pronounced short-term variability in the MR plume during the sampling period, including a retreat and weakening of the surface phytoplankton bloom from May 6 −9. On May 7, the region of highest Chl-*a* extended offshore and encompassed Station 4, whereas by May 9 the bloom had contracted toward the coast, placing Station 4 near the plume margin. This temporal evolution is consistent with the separation of profiles in the T-S diagrams (Fig. 6), where stations sampled on different days fall into distinct hydrographic clusters.

The T-S diagrams (Fig. 6) separated the profiles into four distinct clusters, each representing a different hydrographic regime associated with the MR plume (Fig. 5). The first cluster corresponded to plume-core waters (Sta. 1A-1C), characterized by low surface salinity (< 29 psu), elevated Chl-*a*, and high CDOM and sediment loads in the satellite RGB images. The second cluster represents plume-edge mixing waters (Sta. 3A-3C and Sta. 4C-4D), where intermediate surface salinities (30-32 psu) and strong vertical salinity gradients reflect the interaction between riverine and shelf waters. The third cluster comprises shelf saline waters (Sta. 4A-4B and Sta. 5A-5B), exhibiting higher and more uniform salinity (> 33 psu) and moderate Chl-*a*. The fourth cluster corresponds to offshore oligotrophic waters (Sta. 2), characterized by high salinity, weak stratification, and low satellite-derived Chl-*a*.

Based on the T-S characteristics, plume waters were identified as warm, low-salinity surface waters (S < ∼30-32 psu), whereas shelf waters were cooler,more saline (S ∼ 34-36 psu) and occupied a deeper portion of the water column. At plume-edge stations, the vertical juxtaposition of these two water masses produced a sharp halocline that corresponds to the pycnocline and forms a bottom-trapped saline wedge. A salt wedge is a bottom-trapped intrusion of dense seawater that propagates landward beneath a fresher surface plume, generating a sharp halocline, tilted isohalines, and strong vertical density gradients (Wright & Coleman 1971; Kasai et al. 2010). This structure was clearly expressed in the vertical profiles, where plume waters are confined to the upper ∼10-15 m, overlying intruding shelf waters.

At the coastal station (Sta. 1), the temperature and salinity profiles shared a common structure. A well-defined halocline coincided with steep positive temperature gradients extending to deeper waters (∼ 15 m), a characteristic suggesting the presence of mixed estuarine waters. Profiles 1A and 1B, collected on the same day within a short time interval, show similar water-mass characteristics, whereas profile 1C, measured the next day (approximately 24 hours later), differs in the properties of the surface water mass. Warmer, well-mixed estuarine waters were observed at Sta. 1C.

In contrast, the stations nearest the MR mouth (Stations 4A-4D) showed more pronounced subsurface temperature variations, along with higher surface temperatures than those observed at Sta.1. At Stations 4A-4D, the potential density ranged from 14.5 to 17 kg/m³, with temperature and salinity ranging between ∼23–25 °C and ∼23–25 psu, respectively. Below, a strong halocline separates the estuarine waters from the oceanic waters (S ∼ 35-36 psu; *σ*_*θ*_ = 24-25 kg/m³) at depths > 28 m. Similar stratified profiles were observed at Station 1, although the estuarine water temperatures there were, on average, lower and the salinities higher, respectively.

Given the weak wind waves and tidal currents in this region (Murray 1972), the buoyant MR plume remains at the surface while denser shelf waters intrude landward beneath it due to the strong density contrast (Wright & Coleman 1971). This two-layer circulation maintains a sharp halocline and produces a bottom-trapped saline wedge (Kasai et al. 2010). Stations 3 and 4 exhibited a layer of cooler saltier water below the surface layer.

All profiles at Stations 3 and 5 showed a weaker influence of the river plume (consistent with the satellite images), with surface salinities of ∼ 31 psu. These surface salinities were still lower than those of the deeper waters, but substantially higher than the surface salinities at Stations 1 and 4. The observations at Station 3 pointed to localized stratification at the depth of the thin layers, embedded within a water column that was otherwise actively mixed. This vertical configuration allowed thin layers to persist at the plume–shelf interface while mixing remained strong above and below. At Stations 3A-3C and 4C-4D, a pronounced halocline separated the less saline surface waters (S ∼ 30-32 psu; S ∼ 23.2-25.7 psu) from the denser ocean waters (S ∼ 35-36 psu), forming a salt wedge trapped at the bottom. Thin layers of phytoplankton consistently occured at or immediately above the salinity interface. Overall, the T-S diagrams demonstrated the short-term variability of MR plume dynamics and its variable influence on the sampled stations.

### 3.2. Vertical distribution of plankton

Figure 8 summarizes the vertical profiles of plankton abundance at all stations. Zooplankton concentrations showed no clear relationship with the local thermohaline structure, whereas phytoplankton reached their highest densities in the warm, low-salinity surface layer produced by the MR plume, a feature consistent with nutrient enrichment.

**Fig. 8.**
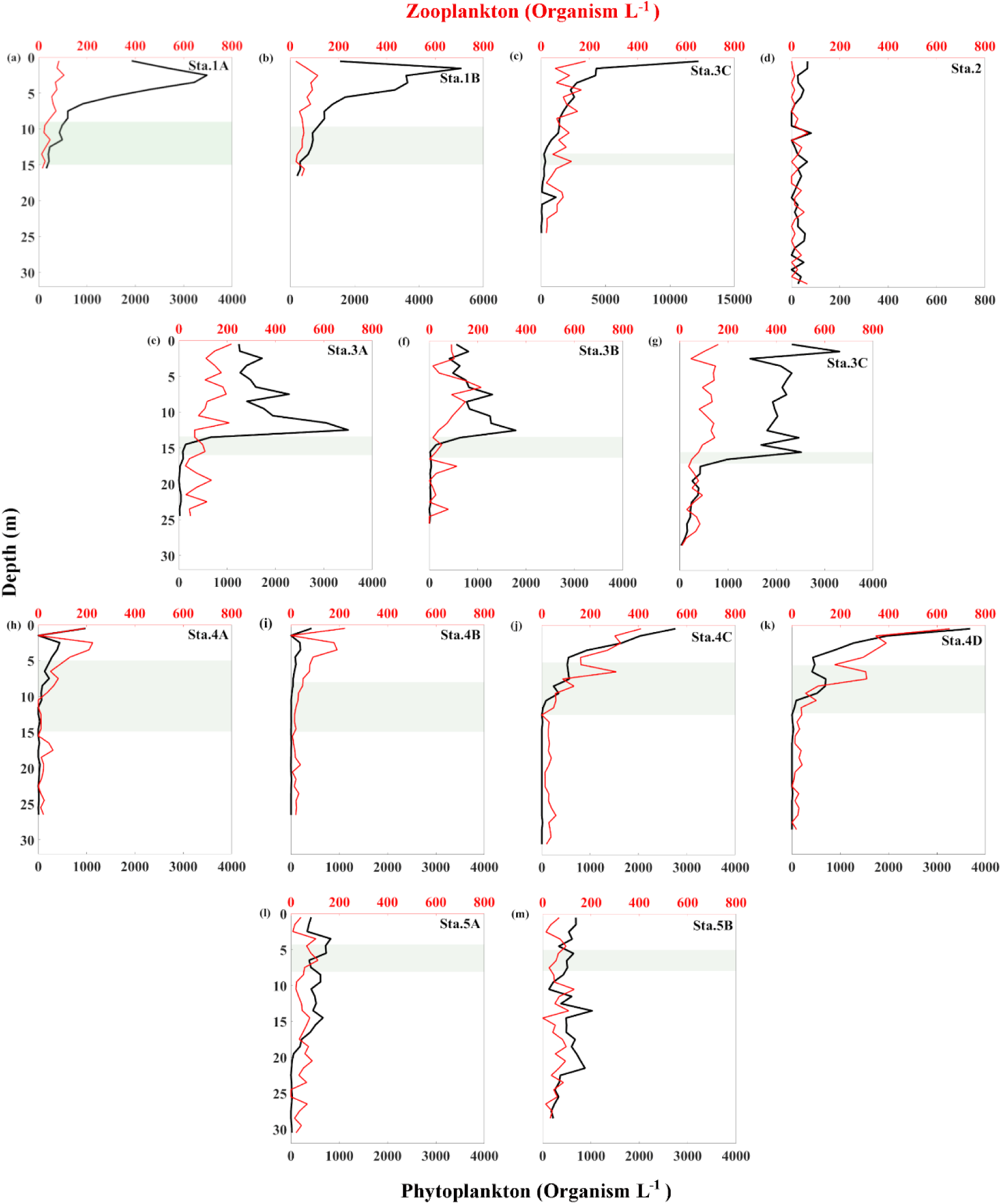
Vertical distribution of total phytoplankton and zooplankton concentrations for all stations visited in the vicinity of the MR plume. The shaded region in each profile indicates the pycnocline region. Note the different horizontal scales within the subplots for the phytoplankton concentration.

Phytoplankton concentrations varied among stations and with depth. At stations 1A-1C, phytoplankton exhibited pronounced surface maxima, with Station 1C showing substantially higher concentrations than 1A and 1B (Fig. 8). At this station, concentrations dropped sharply within the pycnocline and in deeper layers between 11 and 18 m.

In contrast, Stations 2, 4A, 4B, 5A and 5B exhibited consistently lower phytoplankton abundances throughout the water column (Fig. 8). At stations 3A and 3B, plankton concentrations peaked between 10 and 15 m and decreased below this depth, corresponding to the position of the local thermocline. Stations 4C-4D, located at the boundary between the plume and the open ocean, exhibited a monotonic decrease in phytoplankton abundance with depth, with surface maxima above 10 m, with a small secondary maximum around 9 m.

Zooplankton was distributed throughout the entire water column, showing little variation with depth, except for Sta.4, where concentrations peaked above 5 m depth, rapidly decreasing below.

Representative holographic reconstructions of the most frequently encountered phytoplankton and zooplankton taxa in the processed datasets are presented in Figure 9.

**Fig. 9.**
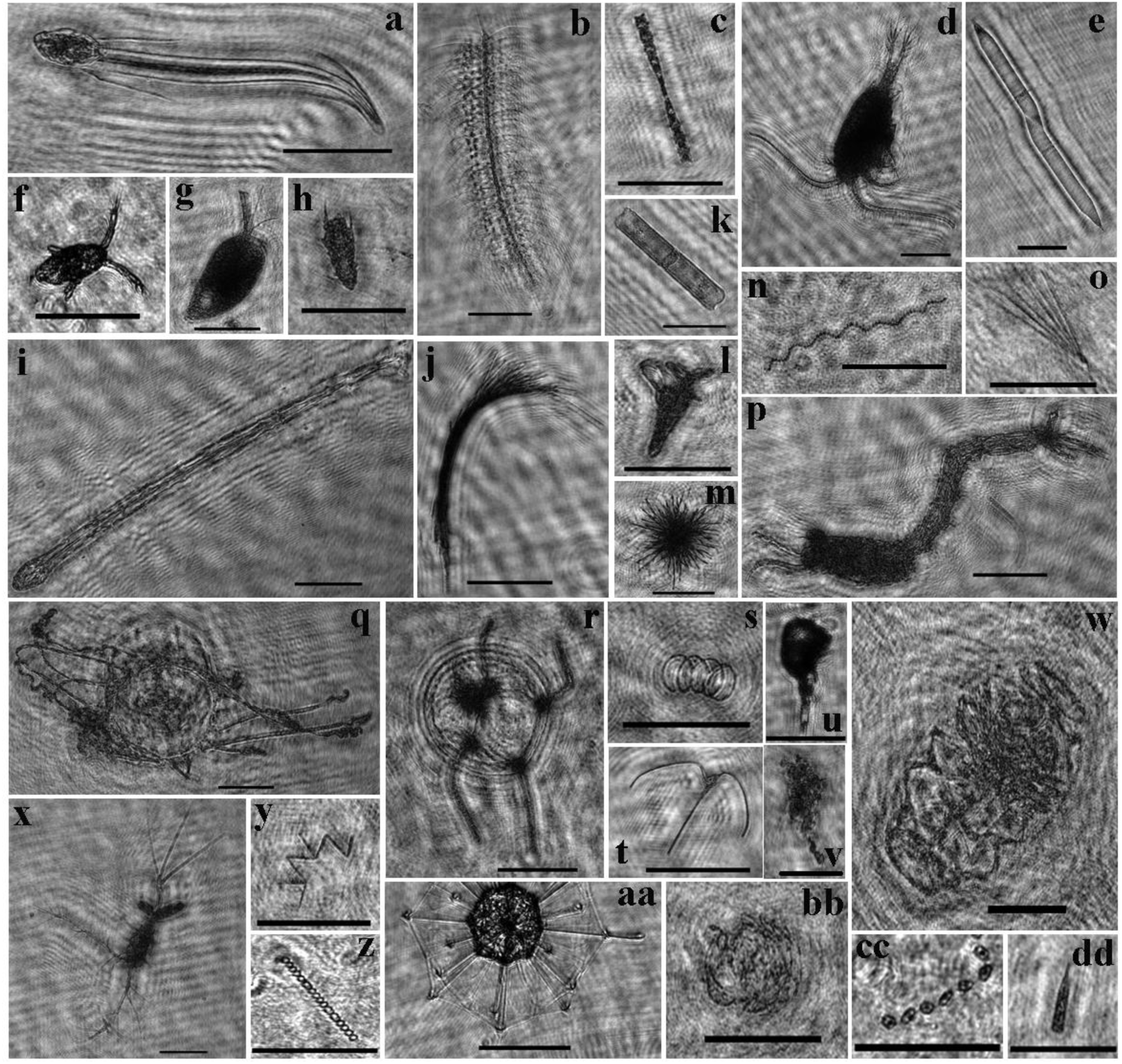
Reconstructed images of some of the most abundant taxa in the study region, obtained after segmentation, reconstruction and automatic classification. The taxonomic identification is denoted by the letters at the top end of each image: a) appendicularian; b) *Chaetoceros* sp.; c) *Helicotheca* sp.; d) copepod (*Temora* sp.); e) *Rhizosolenia* sp.; f) copepod nauplius; g) ostracod; h) polychaete; i) chaetognath; j) *Trichodesmium erythraeum*; k) *Guinardia* sp.; l) pteropod; m) *Trichodesmium thiebautii*; n) *C. debilis*; o) *Lioloma* sp.; p) decapod; q,r) hydromedusa; s) unidentified centric diatom chain; t) *Tripos* sp.; u) copepod (*Corycaeus* sp.); v) aggregate; w) Doliolida; x) *Oithona plumifera*; y) *Thalassionema nitzschioides*; z) *Melosira* sp.; aa) radiozoa; bb) *C. socialis* colonies; cc) *Thalassiosira* sp.; dd) tintinnid. The scale bars (black line) are 1 mm.

### 3.3. Vertical distribution of the major planktonic groups

Chain-forming diatoms dominated the phytoplankton assemblage across the entire depth range at every station, with *Chaetoceros socialis* and *C. debilis* far outnumbering other centric forms (Fig.9 and Fig. 10). Their abundance generally declined from the surface downward, yet clear subsurface maxima emerged: *C. debilis* peaked in the mid-water layer at Stations 3A-3C, whereas *C. socialis* reached comparable maxima at Stations 5A and 5B. The deepest layer at Station 5B sustained dense *C. socialis* chains, averaging 2.5 × 10³ cells L⁻¹. Other commonly observed diatoms included *Chaetoceros curvicetus*, *Thalassionema nitzchioides*, and *Melosira sp.*, followed by chain-forming cyanobacteria (*Trichodesmium* spp.). The *Chaetoceros* category consists of *Chaetoceros curvicetus*, *Chaetoceros coarctatus*, and other unidentified species belonging to the *Chaetoceros* genus (Fig.9b, 9n, 9bb).

**Fig. 10.**
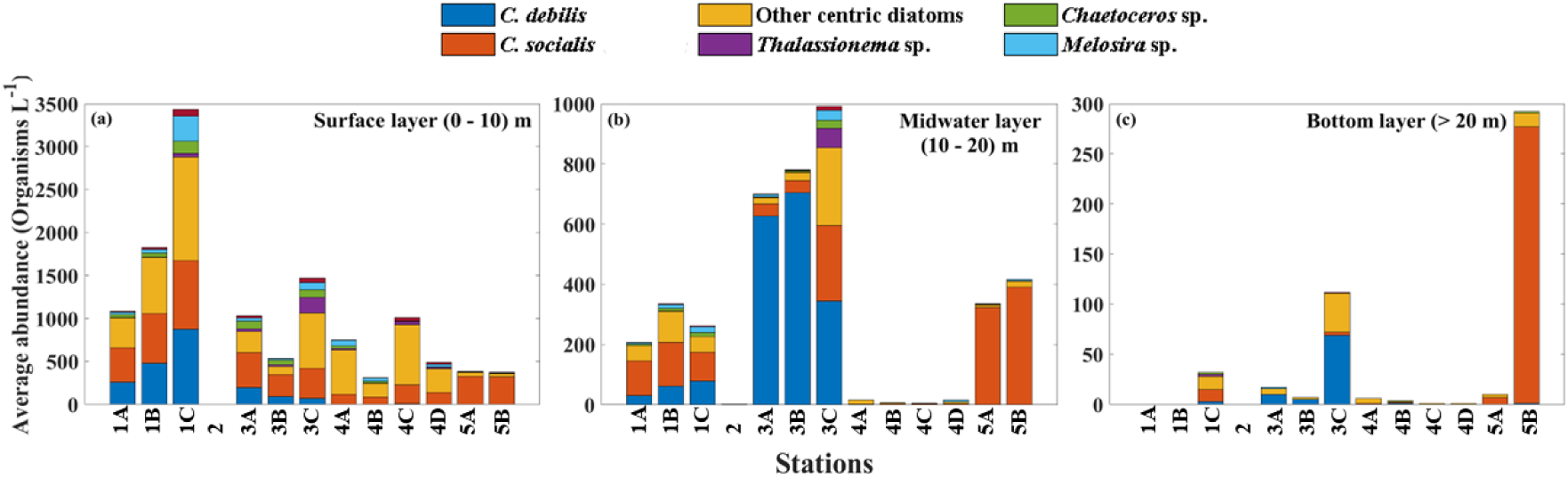
Abundance distribution (Organisms L^-1^) of the most commonly occurring phytoplankton taxa in the analyzed stations (A) surface layer (0–10 m); (B) middle layer (10–20 m) and (C) lower layer (>20 m). Note the different vertical axis limits for each subplot.

Figure 10 showed that Stations 1A-1C and 3A-3C exhibit similar vertical abundance patterns, characterized by maxima within the upper part of the water column, although the absolute phytoplankton abundances differ markedly between these stations. Most organisms were concentrated at depths shallower than 20 m, with a greater tendency for the presence of species such as *C. debilis* and *C. socialis* colonies and other centric diatoms chains. The abundance of these species displayed an increasing trend toward the coast (Sta.1A-1C) and a lower representation in abundance at the intermediate stations Sta.3A-3C. Station 5, located between the coast and the open sea, showed a higher presence of *C. socialis* colonies. Station 2, the only site sampled in deeper waters, exhibited a lower abundance of organisms across all identified taxa (Fig.8 and Fig.10).

Kruskal–Wallis tests revealed significant vertical differences in abundance for all six phytoplankton groups (p ≤ 0.044), indicating strong depth-related structuring (Fig. 10). Dunn’s post-hoc comparisons showed that, for all groups of organisms except *Chaetoceros* sp., the main contrasts occurred between the surface (10 m) and deeper layers (30 m; adjusted p < 0.05), whereas *Chaetoceros* sp. also differed between 20 and 30 m.

A gradient in zooplankton abundances from surface to bottom depth bins was also noted, with a prevalence of appendicularians and calanoid copepods at all stations (Fig. 11). The “ecdysis” category consists of empty crustacean carapaces. Differences in taxonomic composition and concentration between stations 3A and 3B that were taken a few minutes apart may be related to small-scale temporal variations.

**Fig. 11.**
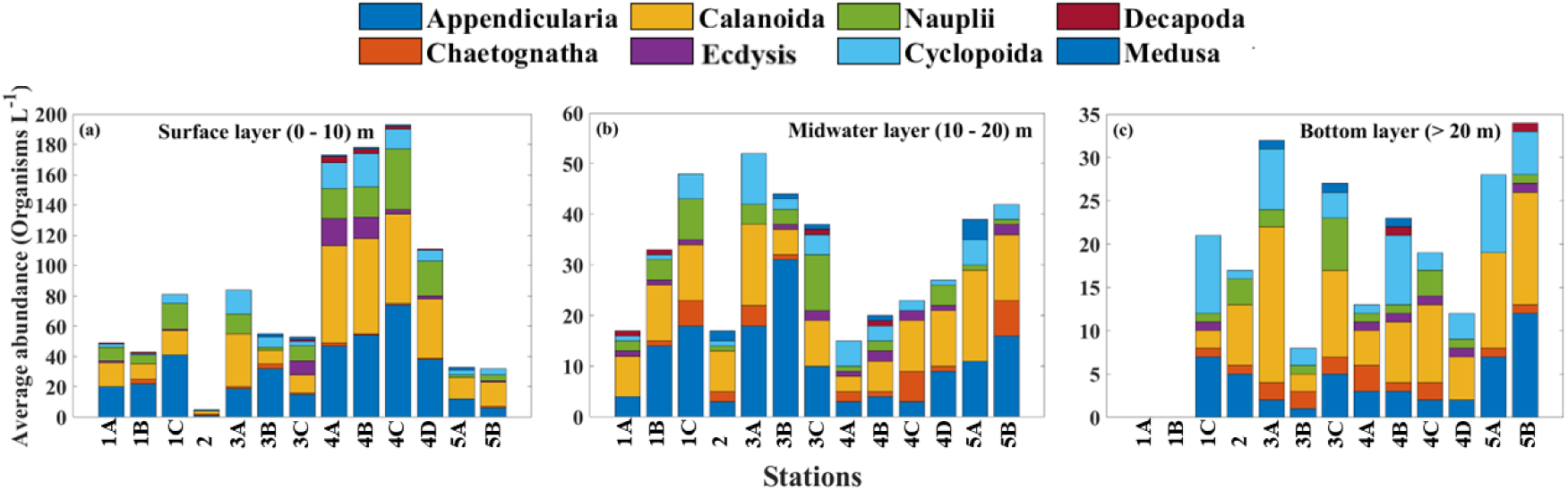
Abundance distribution (Organisms. L^-1^) of the most commonly occurring zooplankton groups in the analyzed stations (A) surface layer (0–10 m); (B) middle layer (10–20 m) and (C) lower layer (>20 m). Note the different vertical axis limits for each subplot.

Significant differences among the three depth layers (df = 2) were detected for appendicularians (H = 19.4, p = 6.1 × 10⁻⁵), copepods (H = 15.9, p = 3.5 × 10⁻⁴), and nauplii (H = 14.1, p = 8.7 × 10⁻⁴). The remaining organism groups did not show significant differences (p > 0.1), whereas ecdysis exhibited a marginally non-significant pattern (H = 5.91, p = 0.052), indicating a potential weak trend of vertical differentiation. Post-hoc comparisons indicated that, in appendicularians and copepods, zooplankton abundance was significantly higher in the upper layer (10 m) than in the deeper layers (20 and 30 m; adjusted p < 0.05), while no significant differences were detected between the middle (20 m) and bottom (>30 m) layers after Bonferroni correction.

### 3.4. Chlorophyll-a and density profiles

Figure 12 displayed profiles of chlorophyll concentration (Chl-*a*) and density (*σ*_*T*_) as a function of water column depth for all analyzed stations.

**Fig. 12.**
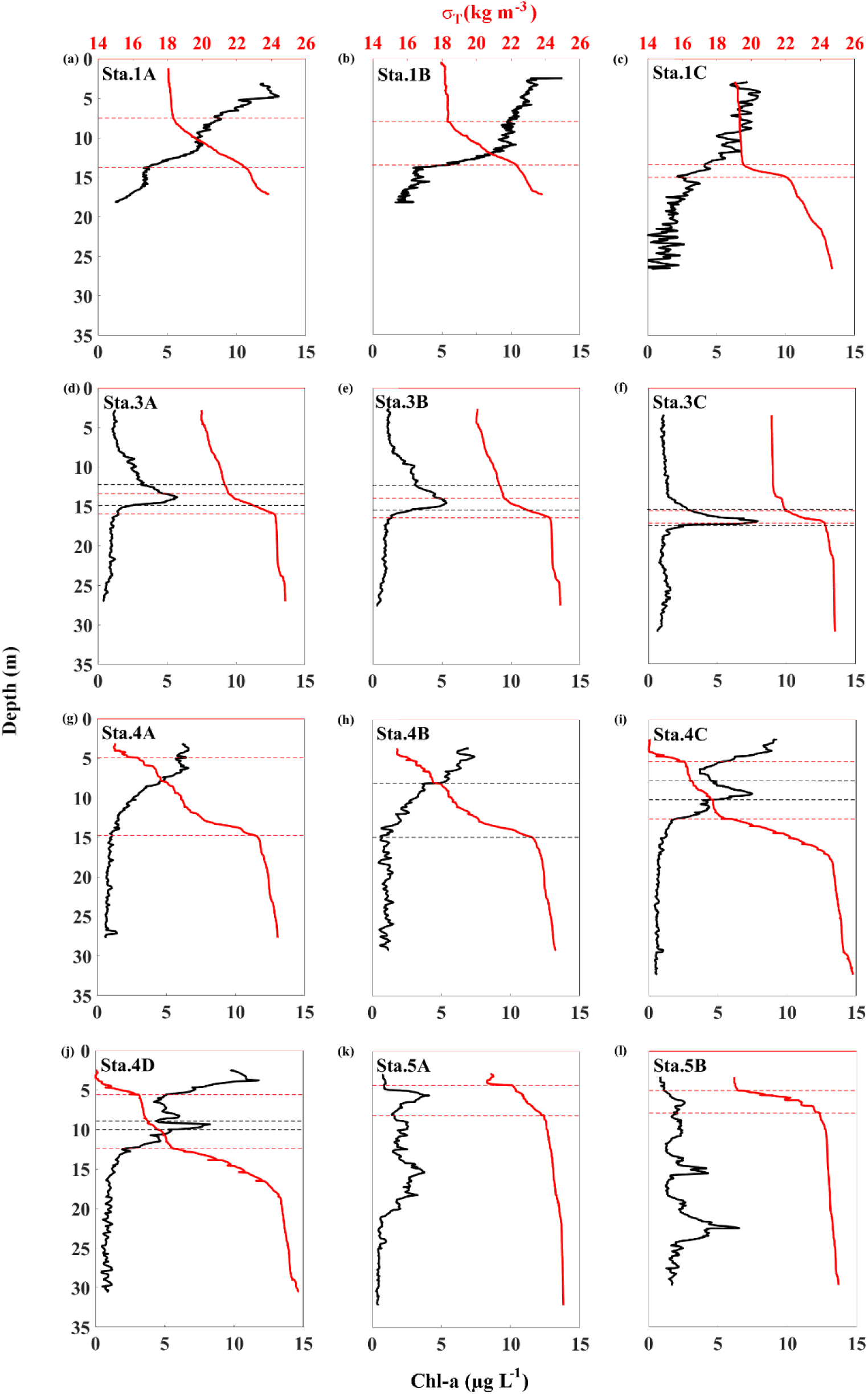
Profiles of chlorophyll a concentration (black) and density *σ*_*T*_ (red), as a function of water column depth, for all analyzed stations. The red dashed lines indicate the pycnocline and the black dashed lines (Sta.3A, 3B, 3C and Sta.4C, 4D) indicate the thin layer thickness. Only the primary pycnocline associated with plume-induced stratification and thin-layer formation is highlighted.

Sta. 3A, 3B and 3C exhibited the formation of a vertically thin, sharply defined pycnocline/halocline at approximately 15 m depth, coinciding with Chl-a peaks. Within the pycnocline at Stations 3A, 3B, and 3C, Chl-*a* peaks of 5.8, 5.4, and 7.9 µg L⁻¹ were observed, coincident with the depths of the thin phytoplankton layers identified in Fig. 8 (black dashed lines in Fig. 12).

In each case, the peak represented a roughly threefold higher concentration compared to the surrounding water column, and its vertical extent remained narrow, i.e., 3.2 m at 3A, 3.5 m at 3B, and 2.2 m at 3C, characteristics that allowed us to identify these structures as thin layers of chlorophyll.

At Station 3A, plankton abundance peaked at 3.6 × 10³ organisms L⁻¹, while at Station 3B the maximum abundance reached 1.8 × 10³ organisms L⁻¹. In both cases, these peaks occurred at depths coincident with the chl-a thin layers identified in the vertical profiles (Fig. 8), as indicated by the black dashed lines in Fig. 12. At station 3C, the water column was more homogeneous: distinct abundance peaks occurred not only at the chlorophyll maximum (2.6 × 10³ organisms L⁻¹) but also at the surface (3.4 × 10³ organisms L⁻¹) (Fig. 8). All three thin layers stood out sharply from secondary chlorophyll peaks elsewhere in the cast, displaying much steeper vertical gradients and a consistent dominance of *Chaetoceros debilis* (Fig. 10).

On May 10, thin layers were observed at Stations 4C and 4D. Chl-*a* reached 7.5 and 8.4 µg L⁻¹, more than triple the background values of 0.6 and 0.9 µg L⁻¹, yet the layers were only 1.5 and 1.25 m thick, respectively (Fig. 12). Phytoplankton concentrations at these depths were modest (∼ 3.4 × 10² and 8.6 × 10² organisms L⁻¹), far below the surface maxima (Fig. 8). The layers contained predominantly *Chaetoceros socialis* chains and other centric diatoms; additionally, a small cohort of appendicularians was present at Station 4C (Figs 9, 10,11).

At Stations 5A and 5B, only minor vertical variability in chl-a was observed, and these features did not satisfy the criteria used to identify thin layers. These maxima, however, did not coincide with the major phytoplankton peaks (Fig. 8).

### 3.5. Redundancy analysis

Redundancy analysis suggested how the hydrographic setting may influence plankton assemblages. Stations 1A-C and 3A-C were located on the positive side of the ordination together with the longest environmental vector (chlorophyll-*a*), indicating that local productivity, rather than temperature or salinity, was likely a primary factor associated with the structure of their plankton assemblages. *C. debilis* and *C. socialis* lay in the same quadrant, supporting the interpretation that these chain-forming diatoms tended to thrive under nutrient-rich, turbid plume conditions.

Axis 2 (14.7%) distinguished stations primarily by stratification intensity. Station 5 (moderately plume-affected) occupied the upper-left quadrant, where elevated salinity and density coincided with moderate chlorophyll concentrations and intermediate plankton abundances. In contrast, the casts from Station 4 were located in the lower-left quadrant, away from the vectors representing the dominant diatom species, suggesting a distinct community configuration not fully captured by the measured hydrographic variables. Although these samples contained relatively few *C. debilis* and *C. socialis* cells, they exhibited a higher contribution from other centric diatoms, consistent with thin layer observations that revealed a secondary chlorophyll maximum dominated by those taxa. Along this axis, lower temperature, salinity, and density were associated with higher numbers of appendicularians, copepod nauplii, and cyclopoid copepods, a pattern most evident at Station 4.

## 4. DISCUSSION

### 4.1. Mechanisms of thin layer formation and persistence

Sub- to mesoscale forcing governs plankton organization by modulating stratification, nutrient delivery, light climate, and turbulence (Acosta-Corridor 2019). Spring winds and the large freshwater–nutrient pulse from the Mississippi-Atchafalaya system combines to generate a buoyant surface plume that caps denser shelf water (Fournier et al. 2016). Where this plume intersects the pycnocline, we observed thin chlorophyll layers at Stations 3A, 3B, and 3C, mirroring the tendency for roughly 70 % of such layers to sit within, or immediately beneath, the density interface (Dekshenieks et al. 2001). The lidar work of Yang et al. (2022) shows that similar thin phytoplankton layers occurring at mid-water depths (10–15 m, 2–6 µg Chl-*a* L⁻¹) are widespread across the plume margin, supporting the idea that our snapshots capture a persistent regional phenomenon.

Various mechanisms have been proposed to elucidate the formation and persistence of thin layers in the oceans (Johnston & Rudnick 2009, Durham & Stocker 2012). Among these, physical stratification associated with strong density gradients and vertical shear has been discussed in previous studies focused on the GoM and elsewhere (Talapatra et al. 2013, Greer et al. 2020). These studies have concluded that thin layers exhibit a tendency to develop in stable areas, where salinity serves as the primary driver of density stratification, typically occurring beneath the mixed layer.

The synoptic satellite observations (Fig. 5) indicate that thin layers developed primarily at stations located at the plume margin rather than within the highly turbid plume core or in offshore oligotrophic waters. This suggests that the transitional zone between riverine and oceanic waters provide optimal conditions for salt wedge formation and enhanced stratification, which in turn favor thin layer development. The co-occurrence of plume-edge conditions in Fig. 5 and strong haloclines in the *in situ* profiles (Fig. 7) support the interpretation that mesoscale plume structure governs the fine-scale vertical organization of phytoplankton in the nGoM.

Previous studies describing frontal systems and plume-shelf interactions provide a physical framework for interpreting our observations (Clayton et al. 2014; Nagai et al. 2012, 2015; Fournier et al. 2016). These studies show that plume margins are characterized by convergent flow, enhanced stratification, and localized shear, conditions that can concentrate plankton and promote the formation of thin layers (Dekshenieks et al. 2001; Greer et al. 2020; Lévy et al. 2018). Our results are consistent with this framework: thin layers were most frequently observed near the plume-shelf interface, where a vertically thin pycnocline persisted despite active mixing above and below. The temporal variability observed at fixed stations further supports the idea that ecosystem responses at plume fronts can evolve rapidly as frontal position and strength change, even when surface salinity remains similar. These findings support the hypothesis that frontal dynamics associated with plume-shelf interactions play a central role in organizing plankton into thin layers in this region.

At Stations 3A and 3B, layer thickness averaged 3.3 m and chlorophyll peaks were threefold above background levels. Twenty-four hours later, Station 3C displayed a thinner (2.2 m) yet more intense layer. Although Station 3C exhibits a pronounced local maximum in buoyancy frequency (N) at the pycnocline (Fig. 13), the stratification above and below this interface is weaker, resulting in reduced depth-averaged (background) stratification of the water column. This contrast among profiles indicates that thin layer evolution is not governed by stratification magnitude alone. Instead, the concurrent cooling below 18 m, together with an indirectly inferred enhancement of vertical shear associated with the compressed pycnocline at the plume–shelf interface, is consistent with an intrusion of denser shelf water that shoaled and compressed the layer toward a local shear maximum, a scenario that can promote shear-driven instability and mixing (Johnston & Rudnick 2009; Greer et al. 2020). However, thinning with increased peak chlorophyll under weaker background stratification is also compatible with biological intensification (e.g., *in situ* growth and/or behavioral accumulation) acting in concert with physical confinement.

**Fig. 13.**
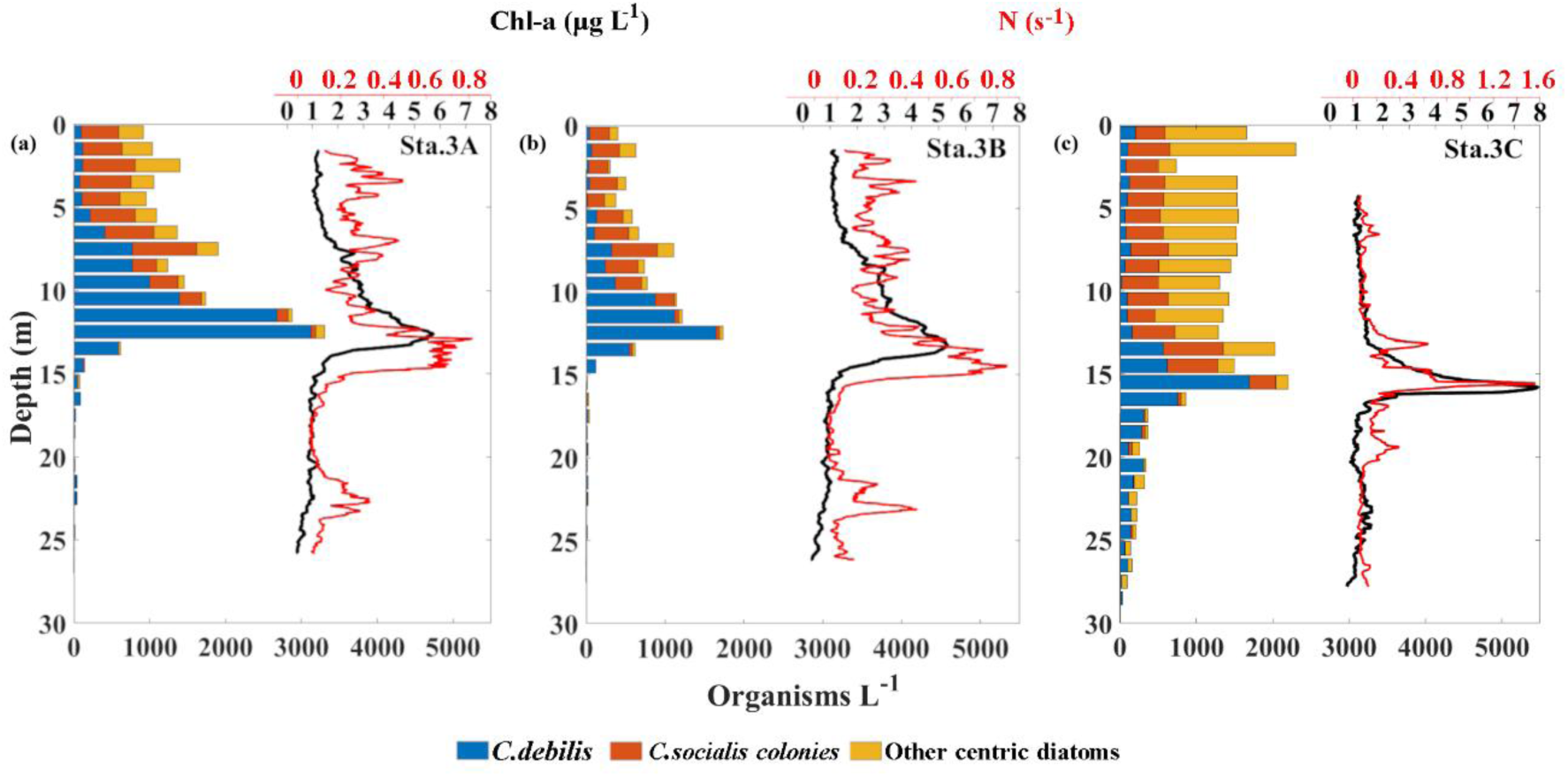
Vertical distribution of dominant phytoplankton taxa for stations 3A, 3B and 3C, where thin layers were detected. Vertical profiles of Chl-*a* (black line; upper axis) and Brunt-Vaisala frequency (red line; upper axis) are also shown.

The combined salinity, density, and chlorophyll profiles (Fig. 7) reveal that the thin layers observed at Stations 3A-3C and 4C-4D are physically linked to the presence of a salt wedge generated by the MR plume. At these stations, a bottom-trapped saline intrusion and a buoyant surface plume generated a compressed pycnocline that acted as a physical trap for phytoplankton cells. The thin layers formed at or just above this interface, where shear and stratification are typically maximized, corroborating the proposed mechanism that salt wedge formation is a key factor in the development of thin layers here. In contrast, when the water column is mixed vertically (e.g., in the surface layer of Station 3C), thin layers are weak or absent.

The satellite imagery (Fig. 5) shows that the surface phytoplankton bloom associated with the MR plume retreated and weakened over only a few days. This rapid change likely contributed to the contrasting hydrographic and biological conditions observed among stations. For example, Station 4 shifted from the bloom core to the plume margin between May 7 and May 9, which helps explain the differences in stratification, salt wedge structure, and thin layer development seen in the *in situ* profiles.

Zooplankton grazing offers an additional pathway for reducing the intensity of the layer, but its influence appeared minimal during our survey. The vertical distributions of appendicularians, nauplii, and calanoids showed only modest increases in abundance near chlorophyll maxima (Figs 11, 14, 15). Benoit-Bird et al. (2010) argued that strong grazer aggregation requires a layer to contain a disproportionately large share of the column-integrated phytoplankton; our layers, although conspicuous, represented < 20 % of total Chl-*a* in the measured portion of the water column, perhaps explaining the weak trophic response. Bochdansky and Bollens (2004) reported that copepods may show a slightly greater tendency to swim through thin diatom layers rather than remain aggregated around them. This behavior does not necessarily imply that copepods did not visit the layer to feed, and the interpretation could change over longer time scales (days to weeks), as variations in river discharge, wind forcing, and shelf circulation modify plume extent, stratification strength, and the persistence of the plume–shelf interface. In our study, the observed thin layers were not analyzed over time, which limits our ability to draw conclusions linking the thinning of these layers to biological factors. Therefore, we attribute the observed thinning primarily to physical rather than biological mechanisms.

### 4.2. Plankton vertical distribution

The T-S diagram (Fig. 6) shows a strong influence of freshwater input from the MR plume in the study area, with a pronounced vertical and lateral mixing zone between plume freshwater and the saline waters of the Gulf at stations 1C, 3B, 4A-4C). Stations 4D, 5A, 5B, located at the outer edge of the plume’s influence, exhibited a reduced freshwater signal. Stations 4A-4D represent repeated profiles collected at the same location over time. While surface salinities remained similar, the vertical structure evolved, with Station 4D showing a weakened plume-driven stratification relative to earlier casts, consistent with temporal variability in plume–shelf interactions. Similar to the dynamics described by Clayton et al. (2014), the transport of distinct water masses can shape ecological patterns across large spatial scales. Nagai and colleagues (e.g., Nagai et al. 2012, 2015) also emphasized how frontal systems and lateral intrusions act as efficient mechanisms for redistributing physical and biochemical properties offshore, influencing plankton community structure through cross-frontal transport and episodic mixing events. These studies highlight that mesoscale and submesoscale processes can promote offshore dispersal of species, enhance nutrient availability, and modify biological productivity far from their source regions.

In Figure 9, a higher presence of *C. debilis* is observed at the station closest to the coast (Sta. 1), shifting to higher occurrences of the same species at slightly greater depths at Sta. 3, where the T-S profile indicated a transition between fresh and saline waters, accompanied by substantial vertical mixing (Sta. 3B and Sta. 3C). Greater vertical mixing at Stations 3B and 3C is inferred from the presence of a deeper, more vertically homogeneous surface layer and weaker temperature and salinity gradients above the pycnocline, while a vertically thin, sharply defined density interface persists at mid-depth. The Kruskal–Wallis test showed that the abundance of *C. debilis* exhibited a significant, although weaker, vertical difference (H = 6.26, df = 2, p = 0.0437) across the three depth layers defined in Fig. 10. In contrast, the remaining taxonomic groups displayed much stronger vertical structuring, with p-values ranging from 1.07 × 10⁻⁴ to 6.1 × 10⁻⁶ (Fig. 10).

At stations located at the outer limits of the plume interaction (Stations 4D, 5A, 5B), *C. debilis* was not detected. This pattern was also observed at Stations 4A and 4B, which showed a stronger river plume influence compared to Station 4C (Fig. 6). Taken together, these results indicate that the observed variation in phytoplankton abundance is not uniform among taxonomic groups, reflecting group-specific responses to environmental conditions, which may be directly related to freshwater transport associated with the river plume, as well as to offshore advection processes similar to those described by Clayton et al. (2014) and Nagai and co-authors (Nagai et al., 2012, 2015). Increased nutrient inputs associated with plume waters, together with enhanced mixing and submesoscale instabilities, may favor fast-growing phytoplankton species or those better adapted to variable environmental conditions (Grover and Chrzanowski, 2004). In every thin chlorophyll layer we observed, the fluorescence peak corresponded to a local enrichment of *C. debilis*, which could imply that this species aggregates more efficiently and thrives in the irradiance-nutrient balance corresponding to the pycnocline.

Although the relatively small water volume imaged in this study is insufficient for rigorous statistics on predator–prey coupling, the depth-resolved trends still provide insights into grazer behavior. Even so, because our quantitative estimates were limited to ROIs ≥ 90 × 90 pixels (>300 µm), picocyanobacteria, small flagellates, and microzooplankton were largely missed, hindering a holistic assessment of plankton community composition and dynamics. This implies that only microalgae in the large microphytoplankton size range were considered here. The selection of larger organisms, together with photoadaptation in response to irradiance, may explain why some phytoplankton peaks do not coincide with the absolute maxima observed in chlorophyll profiles (Fig.8 and Fig.12). Photoadaptation in response to light variations can modify the vertical distribution of phytoplankton in the water column. Under low-light conditions, phytoplankton increase the chlorophyll concentration per cell, thereby intensifying the fluorescence signal detected in measurements; this increase does not necessarily indicate a higher number of cells, but rather a greater pigment content per cell (Moore et al. 2006).

At Station 3, while zooplankton densities remained low, appendicularians and copepods - the principal grazers - formed patchy aggregates throughout the column but tended to avoid the chlorophyll peak, favoring regions with weaker Chl-*a* values at Stations 3A and 3C (Figs 14, 15). Post-hoc comparisons showed that differences occurred mainly between the surface and bottom layers, even after Bonferroni adjustment. Appendicularians exhibited a significant difference in abundance between the surface and bottom layers (pₐ`dj` = 3.19 × 10⁻⁵). Copepods, in turn, showed significant differences between the surface and intermediate layers (pₐ`dj` = 0.0296) and between the surface and bottom layers. Comparisons involving the intermediate layer tended to be non-significant, indicating that this layer often exhibits intermediate abundances and functions as a vertical transition zone.

**Fig. 14.**
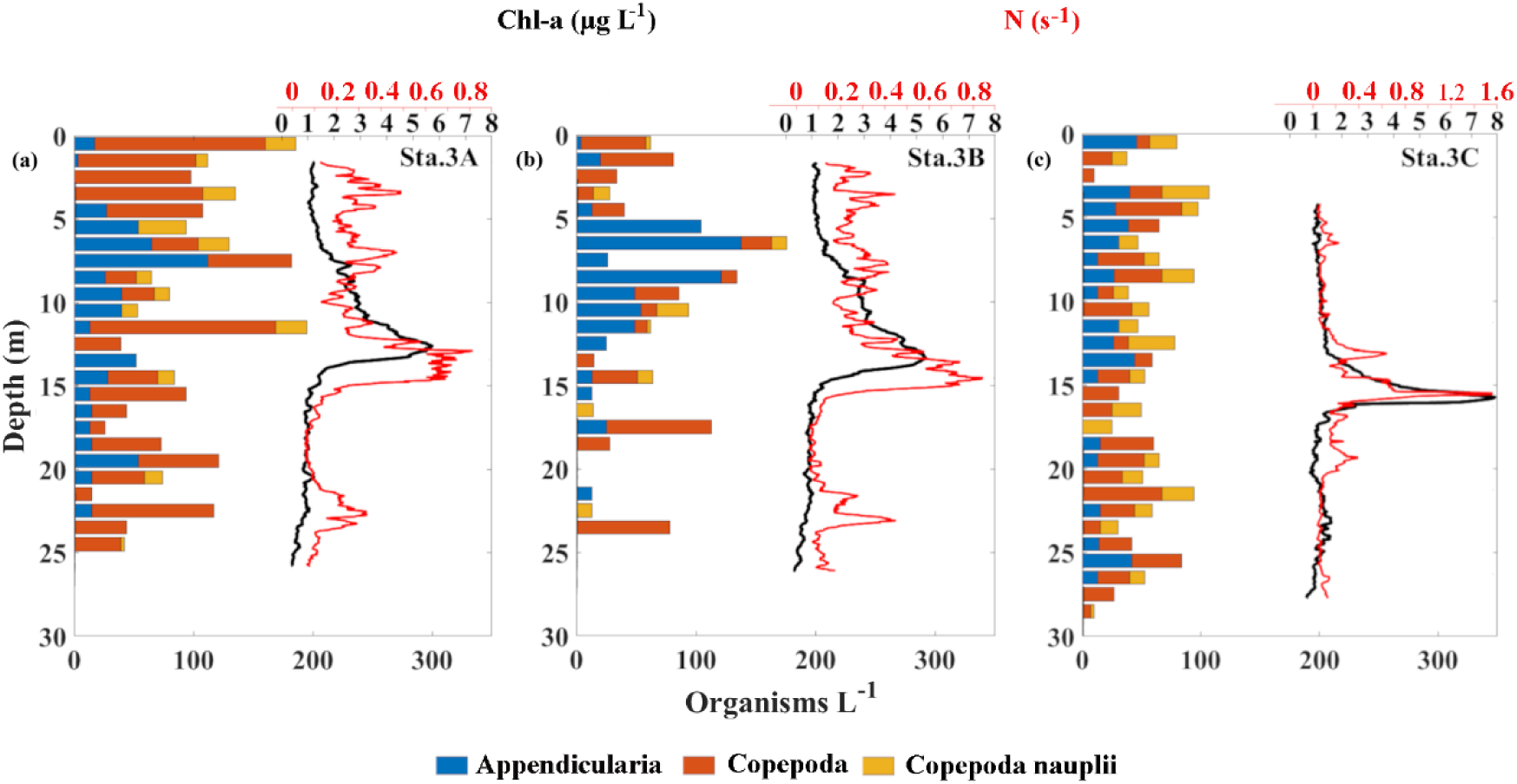
Vertical distribution of dominant zooplankton taxa for stations 3A, 3B and 3C, where thin layers were detected. The same vertical profiles of Chl-*a* (black line; upper axis) and Brunt-Vaisala frequency (red line; upper axis) from Fig. 12 are shown.

**Fig. 15.**
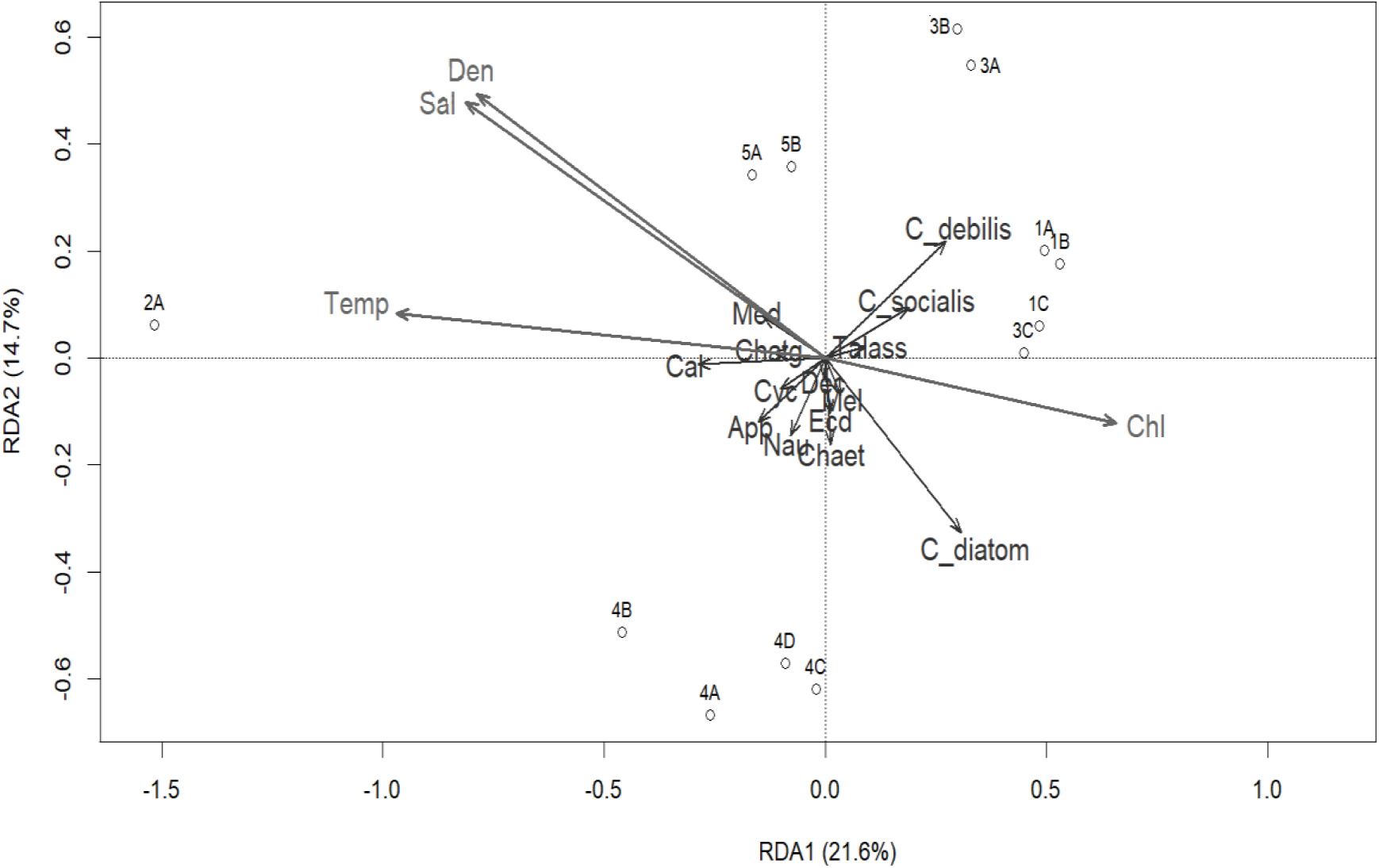
Redundancy-analysis (RDA) biplot showing the relationship between plankton community structure and four hydrographic variables in the northern Gulf of Mexico. Abbreviations are: Talass (*Thalassionema*), Med (Medusa), Chatg (Chaetognatha), Cal (Calanoida), Cyc (Cyclopoida), Dec (Decapoda), Mel (Melosira), Ecd (Ecdise), App (Appendicularia), Nau (Nauplii), Chaet (Chaetoceros, Den (density), Temp (temperature), Sal (salinity).

Only at Station 3B did a modest zooplankton maximum develop just above the layer, driven mostly by appendicularians. Thus, layer formation was decoupled from grazer abundance at this site, at least during daytime, when our profiles were recorded. Consistent with their broad particle size diet (Greer et al. 2013), appendicularians were largely restricted to the upper 12 m even when a thin layer lay beneath them. Such avoidance matches with previous observations of plankton organization at stratified interfaces. High food concentration may be offset by elevated predation risk, sub-optimal oxygen or pH micro-environments, or deterring secondary metabolites, any of which could explain the limited zooplankton interaction with the *C. debilis* layers observed here.

Copepods dominated the zooplankton assemblages, comprising 60.8% of the total zooplankton abundance. The most abundant copepod taxa were nauplii, *Oithona sp.*, and *Oncaea sp*. In contrast, chaetognaths accounted for only 1.81% of the total zooplankton, a low proportion compared to previous studies, where these predatory zooplankton are typically more prevalent and known to tolerate a wide range of environmental conditions (Duro & Saiz 2000). Their habitats include both high-salinity coastal waters (Elliott et al. 2012) and low-salinity layers influenced by mesoscale mixing (McLelland 1989, Gilmartin et al. 2020). These differences in community structure may reflect methodological variability among studies. In particular, the holographic system used here, while offering high-resolution imagery, samples smaller volumes than traditional plankton nets and with a narrow gap between the imaging windows (path length), which may promote avoidance behavior by fast-swimming predators such as chaetognaths, thereby limiting their detection.

Chlorophyll maxima are commonly embedded within meter-scale layers that also concentrate heterotrophic (McManus et al. 2005, Greer et al. 2020, Broullón et al. 2020, McManus et al. 2021, Yang et al. 2022) and mixotrophic plankton (Barua et al. 2024). By compressing primary production into a narrow stratum, these layers can reshape grazer distributions, yet the response is not uniform; toxic or otherwise unpalatable producers often drive zooplankton to congregate just above or below the layer rather than within it (Nielsen et al. 1990, Alldredge et al. 2002).

Copepods and nauplii displayed a heterogeneous, water-column-wide distribution similar to that reported by Talapatra et al. (2013) for the U.S. west coast, where these grazers clustered in layers poor in *Chaetoceros socialis*. Bochdansky and Bollens (2004) also reported the absence of a response by the species *Acartia hudsonica* to diatom aggregations, relating this behavior to the possibility that diatoms are not the preferred food source of copepods. In our profiles, copepods and nauplii exhibited higher abundances in regions with lower relative dominance of *C. debilis*, which may be associated with the presence of other phytoplankton taxa, including *C. socialis* and other centric diatoms. (Figs 13, 14, 15).

Two non-exclusive mechanisms could explain this pattern. First, intense particle concentrations may inhibit copepod feeding by clogging appendages or overwhelming mechanosensory cues, thereby lowering grazing efficiency within the layer. Second, zooplankton may use displacement from the layer as a refuge from visually hunting predators, making brief incursions to feed before retreating to safer depths (Greer et al. 2013). In either scenario, the observed decoupling between phytoplankton and zooplankton highlights the need for simultaneous high-resolution measurements of turbulence, feeding rates, and predator abundance to clarify the behavioral drivers of thin layer utilization.

Field deployments of *in situ* imaging systems now deliver the centimeter-scale detail needed to resolve how thin layer dynamics translate into trophic interactions across entire plankton communities. When the optical record is embedded in a framework of concurrent physical and chemical measurements, it becomes possible to link grazer behavior directly to the evolving hydrographic landscape and to quantify the feedback that regulate patch persistence. Extending such high-resolution sampling over larger spatial and temporal domains will strengthen our capacity to incorporate thin layers - and the behavioral responses they provoke - into predictive ecosystem models and, ultimately, to forecast how changing stratification and nutrient regimes may reshape pelagic food webs.

## 5. CONCLUSIONS

The northern Gulf of Mexico is a physically complex setting where river discharge, wind forcing, and Loop Current eddies interact to create sharp horizontal and vertical gradients. Our high-resolution holographic survey consisting of 13 profiles at 5 different stations in the vicinity of the MR plume demonstrates that this stratification can concentrate certain phytoplankton groups into centimeter- to meter-thick layers located in the pycnocline.

Within these layers, Chl-*a* values tripled relative to background concentrations while the plankton community was dominated by chains of *Chaetoceros debilis* and colonies of *C. socialis*, with cyanobacteria and smaller centric diatoms being secondary contributors. However, because our holographic system captures relatively large particles, in the microplankton size range, future deployments that pair large-volume holography with high-magnification imaging or flow cytometry will be essential for resolving the smallest - but often most abundant - members of the plankton and particle continuum. Copepods, nauplii, and appendicularians formed discrete patches throughout the upper 20 m, but their maxima seldom coincided with the chlorophyll peaks and were frequently displaced into strata of lower chlorophyll, a finding requiring additional investigation on the factors shaping zooplankton grazing in the area.

Because thin layers concentrate biomass and biogeochemical fluxes at small scales, incorporating them into ecosystem models - and into fisheries assessments that rely on accurate prey fields - will require observations with the spatial and temporal fidelity provided by modern *in situ* imaging systems. Extending such measurements across seasons and plume-discharge regimes will allow us to forecast how future shifts in stratification, nutrient supply, and turbulence will propagate through pelagic food webs and comparable river-influenced shelves worldwide.

## Author contributions

Gelaysi Moreno Vega: Conceptualization, data curation, formal analysis, methodology, writing - original draft. Anvita U. Kerkar: Conceptualization, data curation, Writing-original draft, review and editing. Aditya R. Nayak: Resources, Supervision, Writing - Review & Editing. Malcolm McFarland: formal analysis, data curation. Rubens M. Lopes: Resources, Supervision, Writing - Review & Editing.

## Data availability statement

The data is available from the corresponding authors upon reasonable request.

## Acknowledgements

The authors are grateful for the support of Jeffrey Krause of the Dauphin Island Sea Lab and Kanchan Maiti of Louisiana State University for ship time on the R/V Pelican. The authors gratefully acknowledge Laurent Cherubin, Siddhesh Tirodkar and Mingshun Jiang for their useful suggestions on the remote sensing and physical oceanographical data presented in the paper. Data collection efforts were supported by the US National Science Foundation (Awards #1634053 and #1657332). ARN was also supported through a National Academy of Sciences, Engineering, and Medicine (NASEM) Gulf Research Program (GRP) Early Career Fellowship. GMV’s doctoral research was supported by scholarships from National Council for Scientific and Technological Development (CNPq)– Brazil (grant # 141732/2018-0), Coordination for the Improvement of Higher Education Personnel - Brazil (CAPES) - Finance Code 001, and the European Union’s Horizon 2020 research and innovation project “Atlantic Ecosystems Assessment, Forecasting and Sustainability” (AtlantECO) Grant ID: 862923; she later held a postdoctoral position funded by AtlantECO.

## Notes

### Competing Interest Statement

The authors have declared no competing interest.

## LITERATURE CITED

Alldredge AL, Cowles TJ, MacIntyre S, Rines JE, Donaghay PL, Greenlaw CF and others (2002) Occurrence and mechanisms of formation of a dramatic thin layer of marine snow in a shallow Pacific fjord. Mar Ecol Prog Ser 233:1–12

Barua R, Nyman L, Guo B, Johnson MD, Kerkar AU, Hong J and others (2024) *In situ* imaging of a kleptoplastidic ciliate thin layer indicates traditional sampling underestimates oceanic mixotroph biomass. Commun. Earth Environ 5:534

Barua R, Sanborn D, Nyman L, McFarland M, Moore T, Hong J and Nayak AR (2023). *In situ* digital holographic microscopy for rapid detection and monitoring of the harmful dinoflagellate, Karenia brevis. Harmful Algae 123:102401

Benfield MC, Grosjean P, Culverhouse PF, Irigoien X, Sieracki ME, Lopez-Urrutia A and Gorsky G (2007) RAPID: research on automated plankton identification. Oceanogr 20:172–187

Benoit-Bird KJ, Moline MA, Waluk CM and Robbins IC (2010) Integrated measurements of acoustical and optical thin layers I: Vertical scales of association. Cont Shelf Res 30:17–28

Bianchi TS, Filley T, Dria K and Hatcher PG (2004) Temporal variability in sources of dissolved organic carbon in the lower Mississippi River. Geochim Cosmochim Acta 68:959–967

Birch DA, Young WR and Franks PJ (2008) Thin layers of plankton: Formation by shear and death by diffusion. Deep-Sea Res I: Oceanogr Res Pap 55:277–295

Bochdansky AB and Bollens SM (2004) Relevant scales in zooplankton ecology: Distribution, feeding, and reproduction of the copepod Acartia hudsonica in response to thin layers of the diatom Skeletonema costatum. Limnol Oceanogr 49: 625–636

Bochdansky AB, Jericho MH and Herndl GJ (2013) Development and deployment of a point-source digital inline holographic microscope for the study of plankton and particles to a depth of 6000 m. Limnol Oceanogr: Methods 11:28–40

Broullón E, López-Mozos M, Reguera B, Chouciño P, Doval MD, Fernández-Castro B and others (2020) Thin layers of phytoplankton and harmful algae events in a coastal upwelling system. Prog Oceanogr 189:102449

Brown OB, Minnett PJ, Evans R, Kearns E, Kilpatrick K, Kumar A,… and Závody A (1999) MODIS infrared sea surface temperature algorithm algorithm theoretical basis document version 2.0. University of Miami, 31, 098–33

Churnside JH and Donaghay PL (2009) Thin scattering layers observed by airborne lidar. ICES J Mar Sci 66:778–789

Clayton S, Nagai T and Follows MJ (2014) Fine scale phytoplankton community structure across the Kuroshio Front. J Plankton Res 36: 1017–1030

Cowles TJ, Desiderio RA and Carr ME (1998) Small-scale planktonic structure: persistence and trophic consequences. Oceanogr 11:4–9

Dagg M, Sato R, Liu H, Bianchi TS, Green R and Powell R (2008) Microbial food web contributions to bottom water hypoxia in the northern Gulf of Mexico. Cont Shelf Res 28:1127–1137

Dagg MJ and Breed GA (2003) Biological effects of Mississippi River nitrogen on the northern Gulf of Mexico—a review and synthesis. J Mar Syst 43: 133–152

da Silva CE and Castelao RM (2018) Mississippi River plume variability in the Gulf of Mexico from SMAP and MODIS-Aqua observations. J Geophys Res Oceans 123: 6620–6638

Dekshenieks MM, Donaghay PL, Sullivan JM, Rines JE, Osborn TR and Twardowski MS (2001) Temporal and spatial occurrence of thin phytoplankton layers in relation to physical processes. Mar Ecol Prog Ser 223:61–71

Donaghay PL, Rines HM and Sieburth JM (1992) Simultaneous sampling of fine scale biological, chemical, and physical structure in stratified waters. Ergebn Limnol *ERLIA* 6:36

Dougherty ER and Lotufo RA (2003) Hands-on morphological image processing (Vol. 59). SPIE press.

Durham WM, Kessler JO and Stocker R (2009) Disruption of vertical motility by shear triggers formation of thin phytoplankton layers. Science 323:1067–1070

Durham WM and Stocker R (2012) Thin phytoplankton layers: characteristics, mechanisms, and consequences. Annu Rev Mar Sci 4:177–207

Dyomin VV, Polovtsev IG and Davydova AY (2017, November) Fast recognition of marine particles in underwater digital holography. In 23rd International Symposium on Atmospheric and Ocean Optics: Atmospheric Physics p 467–471. SPIE.

Dyomin V, Davydov A, Morgalev S, Kirillov N, Olshukov A, Polovtsev I and Davydov S (2020) Monitoring of plankton spatial and temporal characteristics with the use of a submersible digital holographic camera. F Mar Sci 7:653

Elliott DT, Pierson JJ and Roman MR (2012). Relationship between environmental conditions and zooplankton community structure during summer hypoxia in the northern Gulf of Mexico. J Plankton Res 34:602–613

Fahnenstiel GL, McCormick MJ, Lang GA, Redalje DG, Lohrenz SE, Markowitz M, and others (1995) Taxon-specific growth and loss rates for dominant phytoplankton populations from the northern Gulf of Mexico. Mar Ecol Prog Ser. Oldendorf 117:229–239

Fournier S, Lee T and Gierach MM (2016) Seasonal and interannual variations of sea surface salinity associated with the Mississippi River plume observed by SMOS and Aquarius. Remote Sens Environ 180:431–439

Fournier S, Reager JT, Dzwonkowski B and Vazquez-Cuervo J (2019) Statistical mapping of freshwater origin and fate signatures as land/ocean “regions of influence” in the Gulf of Mexico. J Geophys Res Oceans 124: 4954–4973

Franks PJ (1995) Thin layers of phytoplankton: a model of formation by near-inertial wave shear. Deep-Sea Res I: Oceanogr Res Pap 42:75–91

Gierach MM, Vazquez-Cuervo J, Lee T and Tsontos VM (2013) Aquarius and SMOS detect effects of an extreme Mississippi River flooding event in the Gulf of Mexico. Geophys Res Lett 40: 5188–5193

Gilmartin J, Yang Q and Liu H (2020) Seasonal abundance and distribution of chaetognaths in the northern Gulf of Mexico: The effects of the Loop Current and Mississippi River plume. Cont Shelf Res 203:104146

Goulart AJH, Morimitsu A, Jacomassi R, Hirata N and Lopes R (2021) Deep learning and t-SNE projection for plankton images clusterization. In OCEANS 2021: San Diego–Porto p 1–4. IEEE.

Greer AT, Boyette AD, Cruz VJ, Cambazoglu MK, Dzwonkowski B, Chiaverano LM and others (2020) Contrasting fine-scale distributional patterns of zooplankton driven by the formation of a diatom-dominated thin layer. Limnol Oceanogr 65:2236–2258

Greer AT, Cowen RK, Guigand CM, McManus MA, Sevadjian JC and Timmerman AH (2013) Relationships between phytoplankton thin layers and the fine-scale vertical distributions of two trophic levels of zooplankton. J Plankton Res 35:939–956

Grover JP and Chrzanowski TH (2004) Limiting resources, disturbance, and diversity in phytoplankton communities. Ecol Monogr 74: 533–551

Hernández-Hernández N, Santana-Falcón Y, Estrada-Allis S and Arístegui J (2021) Short-term spatiotemporal variability in picoplankton induced by a submesoscale front south of gran Canaria (Canary Islands). Front Mar Sci 8:592703

Holliday DV, Pieper RE, Greenlaw CF and Dawson JK (1998) Acoustical sensing of small-scale vertical structures in zooplankton assemblages. Oceanogr 11:18–23

Holliday DV, Donaghay PL, Greenlaw CF, McGehee DE, McManus MM, Sullivan JM and Miksis JL (2003) Advances in defining fine-and micro-scale pattern in marine plankton. Aquat Living Resou 16:131–136

Hu C, Lee Z and Franz B (2012) Chlorophyll aalgorithms for oligotrophic oceans: A novel approach based on three-band reflectance difference. J Geophys Res Oceans: 117(C1).

Johnston TS and Rudnick DL (2009) Observations of the transition layer. J Phys Oceanogr 39:780–797

Kasai A, Kurikawa Y, Ueno M, Robert D and Yamashita Y (2010) Salt-wedge intrusion of seawater and its implication for phytoplankton dynamics in the Yura Estuary, Japan. Estuar Coast Shelf Sci 86: 408–414

Koukaras K and Georgios N (2004) Dinophysis blooms in Greek coastal waters (Thermaikos Gulf, NW Aegean Sea). J Plankton Res 26.4:445–457

Legendre P and Anderson MJ (1999) Distance-based redundancy analysis: testing multispecies responses in multifactorial ecological experiments. Ecol Monogr 69:1–24

Lévy M, Franks PJ and Smith KS (2018) The role of submesoscale currents in structuring marine ecosystems. Nat Commun 9:4758

Liu H, Gilmartin J, Sluis MZ, Kobari T, Rooker J, Bi H and Quigg A (2024) Dynamic oceanographic influences on zooplankton communities over the northern Gulf of Mexico continental shelf. J Sea Res 199: 102501

Lohrenz SE, Dagg MJ, Whitledge TE (1990) Enhanced primary production at the plume/oceanic interface of the Mississippi River. Cont Shelf Res 10: 639–64

Lohrenz SE, Fahnenstiel GL, Redalje DG, Lang GA, Dagg MJ, Whitledge TE, Dortch Q (1999) Nutrients, irradiance, and mixing as factors regulating primary production in coastal waters impacted by the Mississippi River plume. Cont Shelf Res 19: 1113–41

Lohrenz SE, Redalje DG, Cai WJ, Acker J and Dagg M (2008) A retrospective analysis of nutrients and phytoplankton productivity in the Mississippi River plume. Cont Shelf Res 28:1466–1475

Lotufo RA, Audigier R, Saúde AV and Machado RC (2023) Morphological image processing. In Microscope image processing. Academic Press p 75–117.

McLelland JA (1989) An illustrated key to the Chaetognatha of the northern Gulf of Mexico with notes on their distribution. Gulf Caribb Res 8:145–172

McManus MA, Greer AT, Timmerman AH, Sevadjian JC, Woodson CB, Cowen R and others (2021) Characterization of the biological, physical, and chemical properties of a toxic thin layer in a temperate marine system. Mar Ecol Prog Ser 678:17–35

McManus MA, Alldredge AL, Barnard AH, Boss E, Case JF, Cowles TJ and others (2003) Characteristics, distribution and persistence of thin layers over a 48hour period. Mar Ecol Prog Ser 261:1–19

McManus MA, Cheriton OM, Drake PT, Holliday DV, Storlazzi CD, Donaghay PL and others (2005) Effects of physical processes on structure and transport of thin zooplankton layers in the coastal ocean. Mar Ecol Prog Ser 301:199–215

McManus MA, Sevadjian JC, Benoit-Bird KJ, Cheriton OM, Timmerman AH and Waluk CM (2012) Observations of thin layers in coastal Hawaiian waters. Estuaries coasts 35:1119–1127

Moore CM., Suggett DJ, Hickman AE, Kim YN, Tweddle JF, Sharples J,… and Holligan PM (2006). Phytoplankton photoacclimation and photoadaptation in response to environmental gradients in a shelf sea. Limnol Oceanogr 51:936–949

Moreno G, Ascaneo JS, Ricardo JO, Leandro T, Arias Y, Strickler JR, Lopes RM (2020) A new focus detection criterion in holograms of planktonic organisms. Pattern Recognit Lett 138:497–506

Murray SP (1972). Observations on wind, tidal, and density-driven currents in the vicinity of the Mississippi River delta. In: Shelf Sediment Transport: Process and Pattern.

Nagai T, Inoue R, Tandon A and Yamazaki H (2015) Evidence of enhanced double-diffusive convection below the main stream of the K uroshio E xtension. J Geophys Res: Oceans 120: 8402–8421

Nagai T, Tandon A, Yamazaki H, Doubell MJ and Gallager S (2012) Direct observations of microscale turbulence and thermohaline structure in the Kuroshio Front. J Geophys Res Oceans 117(C8).

Nardelli SC and Twardowski MS (2016) Assessing the link between chlorophyll concentration and absorption line height at 676 nm over a broad range of water types. Opt Express 24: A1374–A1389

Nayak AR, McFarland MN, Sullivan JM and Twardowski MS (2018a) Evidence for ubiquitous preferential particle orientation in representative oceanic shear flows. Limnol Oceanogr 63:122–143

Nayak AR, McFarland MN, Twardowski MS and Sullivan JM (2018b) On plankton distributions and biophysical interactions in diverse coastal and limnological environments. In SPIE Ocean Sensing and Monitoring X 10631:204–215

Nayak AR, Malkiel E, McFarland MN, Twardowski MS and Sullivan JM (2021) A review of holography in the aquatic sciences: *in situ* characterization of particles, plankton, and small-scale biophysical interactions. Front Mar Sci 7:572147

O’Connor BS, Muller-Karger FE, Nero RW, Hu C and Peebles EB (2016) The role of Mississippi River discharge in offshore phytoplankton blooming in the northeastern Gulf of Mexico during August 2010. Remote Sens Environ 173: 133–144

Oksanen J, Blanchet FG, Friendly M, Kindt R, Legendre P, McGlinn D and Wagner H (2019) vegan: Community Ecology Package. R package version 2.5–5

O’Reilly JE, Maritorena S, Mitchell BG, Siegel DA, Carder KL, Garver SA,… and McClain C (1998) Ocean color chlorophyll algorithms for SeaWiFS. J Geophys Res Oceans 103: 24937–24953

Pedersen TL (2019) The composer of plots CRAN: Contributed Packages 10.32614/CRAN.package.patchwork.

Penninck SB, Lopes RM, Lima JF and McManus MA (2021) Thin layers in the coastal zone of Ubatuba, Brazil: Mechanisms of formation and dissipation. Limnol Oceanogr 66:558–574

Pond S and Pickard GL (1983) Introductory dynamical oceanography. Gulf Professional Publishing

R Core Team. (2024). *R*: A language and environment for statistical computing. R Foundation for Statistical Computing. https://www.R-project.org/

Scavia D, Rabalais NN, Turner RE, Justić D, Wiseman Jr WJ (2003) Predicting the response of Gulf of Mexico hypoxia to variations in Mississippi River nitrogen load. Limnol Oceanogr 48:951–6

Schiller RV, Kourafalou VH, Hogan P and Walker ND (2011) The dynamics of the Mississippi River plume: Impact of topography, wind and offshore forcing on the fate of plume waters. J Geophys Res: Oceans 116(C6)

Selph K, Swalethorp R, Stukel M, Kelly T, Knapp A, Fleming K and others (2022) Phytoplankton community composition and biomass in the oligotrophic Gulf of Mexico. J Plankton Res 44:618–637

Sigman DM and Hain MP (2012) The biological productivity of the ocean. Nat Sci Educ 3:21

Sullivan JM, Twardowski MS, Donaghay PL and Freeman SA (2005) Use of optical scattering to discriminate particle types in coastal waters. Appl Opt 44:1667–1680

Sullivan JM, Donaghay PL and Rines JE (2010) Coastal thin layer dynamics: consequences to biology and optics. Cont Shelf Res 30:50–65

Talapatra S, Hong J, McFarland M, Nayak AR, Zhang C, Katz J and others (2013) Characterization of biophysical interactions in the water column using *in situ* digital holography. Mar Ecol Prog Ser 473:29–51

Tan S and Wang S (2013) An approach for sensing marine plankton using digital holographic imaging. Optik 124:6611–6614

Taylor AH, Harris JRW and Aiken J (1986) The interaction of physical and biological processes in a model of the vertical distribution of phytoplankton under stratification. In Elsevier oceanography series 42:313–330

Turner RE, Rabalais NN (1994) Coastal eutrophication near the Mississippi river delta. Nature 368: 619–21

Wawrik B and Paul JH (2004) Phytoplankton community structure and productivity along the axis of the Mississippi River plume in oligotrophic Gulf of Mexico waters. AME 35:185–196

Weng J, Zhong J, Hu C (2008) Digital reconstruction based on angular spectrum diffraction with the ridge of wavelet transform in holographic phase-contrast microscopy. Opt. Express 16:21971–21981

Wickham H (2011) Ggplot2. WIREs Computational Statistics 3: 180–185. 10.1002/wics.147

Wickham H, Averick M, Bryan J, Chang W, McGowan L, François R, Grolemund G, Hayes A, Henry L, Hester J, Kuhn M, Pedersen T, Miller E, Bache S, Müller K, Ooms J, Robinson D, Seidel D, Spinu V,… Yutani H (2019) Welcome to the Tidyverse. Journal of OpenSource Software 4: 1686. 10.21105/joss.01686

Wright LD and Coleman JM (1971) Effluent expansion and interfacial mixing in the presence of a salt wedge, Mississippi River delta. J Geophys Res 76:8649–8661

Yang Y, Pan H, Zheng D, Zhao H, Zhou Y and Liu D (2022) Characteristics and formation conditions of thin phytoplankton layers in the northern Gulf of Mexico revealed by airborne lidar. Remote Sens 14:4179

